# tRNAscan-SE 2.0: Improved Detection and Functional Classification of Transfer RNA Genes

**DOI:** 10.1101/614032

**Authors:** Patricia P. Chan, Brian Y. Lin, Allysia J. Mak, Todd M. Lowe

**Author notes:** Joint Authors (Patricia P. Chan, Brian Y. Lin).

## Abstract

tRNAscan-SE has been widely used for transfer RNA (tRNA) gene prediction for over twenty years, developed just as the first genomes were decoded. With the massive increase in quantity and phylogenetic diversity of genomes, the accurate detection and functional prediction of tRNAs has become more challenging. Utilizing a vastly larger training set, we created nearly one hundred specialized isotype-and clade-specific models, greatly improving tRNAscan-SE’s ability to identify and classify both typical and atypical tRNAs. We employ a new comparative multi-model strategy where predicted tRNAs are scored against a full set of isotype-specific covariance models, allowing functional prediction based on both the anticodon and the highest-scoring isotype model. Comparative model scoring has also enhanced the program’s ability to detect tRNA-derived SINEs and other likely pseudogenes. For the first time, tRNAscan-SE also includes fast and highly accurate detection of mitochondrial tRNAs using newly developed models. Overall, tRNA detection sensitivity and specificity is improved for all isotypes, particularly those utilizing specialized models for selenocysteine and the three subtypes of tRNA genes encoding a CAU anticodon. These enhancements will provide researchers with more accurate and detailed tRNA annotation for a wider variety of tRNAs, and may direct attention to tRNAs with novel traits.

## INTRODUCTION

Transfer RNAs (tRNAs) are ubiquitous in all living organisms as the key translator of the nucleic acid code into proteins. tRNAscan-SE (1) is the most widely employed tool for identifying and annotating tRNA genes in genomes. With over nine thousand citations, its users include sequencing centers, database annotators, RNA biologists, and other researchers. To increase the ease of use for scientists who cannot install and utilize UNIX-based software, the tRNAscan-SE On-line website (2,3) provides quick, in-depth tRNA analysis. tRNAs predicted using tRNAscan-SE are also available in the Genomic tRNA Database (GtRNAdb) (4,5) for thousands of genomes, enabling the research community to browse high-quality tRNA collections across all three domains of life.

Numerous other tRNA detection and classification methods have become available since tRNAscan-SE was first developed, including ARAGORN (6), which detects both tRNA genes and tmRNA genes; DOGMA (7), ARWEN (8), and MiTFi (9), which are designed for annotating tRNAs in various types of organellar genomes; TFAM (10), which classifies bacterial tRNAs based on log-odds profiles built from covariance models; tRNAfinder (11), a rule-based program that detects tRNAs through secondary structure; and SPLITS (12), which aims to discover split and intron-containing tRNAs primarily in archaeal genomes. All of these methods were designed to either improve upon or complement tRNAscan-SE, and notably, many depend on tRNAscan-SE’s core detection software. None of these programs have been improved upon for roughly a decade or more.

The original tRNAscan-SE implementation pioneered the large-scale use of covariance models (CMs) (13) to annotate RNA genes in genomes, predating the invaluable Rfam database (14). By training on structurally aligned sequences of an RNA family, CMs capture RNA conservation via stochastic context-free grammars that integrate both primary sequence and secondary structure information (13,15). Searches using a CM trained on an alignment of 1,415 tRNAs extracted from the Sprinzl tRNA database (16,17) yielded unparalleled sensitivity and specificity (13). However, these gold-standard searches were too slow to scale up to scan large genomes – at just 20 nucleotides per second on contemporary computers, it would have taken almost ten CPU-years to scan the human genome (1). To speed up CM-based tRNA searches to meet the demands of full genome annotation, the original tRNAscan-SE used two fast and sensitive algorithms as first-pass screens to identify putative tRNAs (18,19). When used in combination with CM searches, this strategy increased the overall speed by several orders of magnitude. However, this also greatly restricted algorithmic flexibility, making detection of specialized classes of tRNAs, such as mitochondrial tRNAs, more difficult.

With the innovation of a second-generation CM search algorithm that is optimized and much faster (20), tRNAscan-SE’s first-pass scanners can now be eliminated, vastly simplifying improvements. Adding new, specialized tRNA search capabilities and better functional classification is now only limited by curation of the high-quality alignments needed for CM model creation. tRNAscan-SE 2.0 addresses many limitations in the former version by leveraging new research in tRNA function and by deploying a large suite of new CMs derived from the massively expanded universe of tRNA genes uncovered in the genome era.

Here we describe the most significant update of tRNAscan-SE since its initial release to demonstrate (1) improved sensitivity and performance via incorporation of Infernal 1.1 covariance model search software (20); (2) updated and expanded search models leveraging a more representative diversity of tRNA genes from thousands of newly sequenced genomes; (3) better functional classification of tRNAs, based on comparative analysis of isotype-specific tRNA CMs, (4) accurate identification of vertebrate mitochondrial tRNAs, (5) the ability to detect multiple and/or noncanonically positioned introns in archaeal tRNAs, and (6) a new “high confidence” filter to better discriminate tRNA-derived repetitive elements from canonical tRNAs. With these new capabilities and enhancements, we show that tRNAscan-SE 2.0 maintains the original program’s high sensitivity and selectivity at essentially no cost in execution efficiency.

## MATERIAL AND METHODS

### Genomes analyzed and availability of tRNAscan-SE prediction results

Predicted tRNA genes in nuclear genomes using tRNAscan-SE 2.0 are currently available to browse in the Genomic tRNA Database (GtRNAdb) (5) as well as by file download. The genome assemblies used in this study include 4,036 bacteria, 217 archaea, and 540 eukaryotes. Although these are not exhaustive analyses of all available genomic sequences (which are added constantly), it constitutes a good representation of high quality, substantially complete genomes. Bacterial, archaeal and fungal genomes were obtained from NCBI GenBank (21) while the other eukaryotic genomes were obtained from the UCSC Genome Browser (22), NCBI GenBank (21), and JGI Phytozome (23). For evaluating the mitochondrial tRNA predictions in vertebrates, 3,345 mitochondrial genomes were retrieved from NCBI RefSeq (24). The specific genomes with tRNA sets presented in this manuscript are listed in Supplementary Table S1.

### tRNA search modes

In default search mode, the new tRNAscan-SE 2.0 uses Infernal 1.1 (20) as the state-of-the-art sequence search engine to find and score tRNA genes (Figure 1). The original tRNAscan-SE v1 used tRNAscan (18) and software implementing an algorithm from Pavesi and colleagues (19) as the sensitive first-pass candidate-gathering searches, and Infernal’s forerunner, COVE (13), as the high-specificity tRNA detector. This v1 search mode is still available in 2.0 as the “legacy mode” (-L) for researchers who wish to make backward version comparisons. The Infernal software implements profile stochastic context-free grammars (SCFGs), also known as covariance models because of their ability to detect covariation in conserved RNA secondary structures. Covariance models can be created to identify members of any RNA gene family based on structurally aligned, trusted examples which serve as training sets. tRNAscan-SE 2.0 (Figure 1) employs a combination of 76 different covariance models (Table 1, Supplementary Figure S1) for identifying and classifying the many different types and biological sources of tRNAs in two phases. In step one, models trained on all tRNAs from species in each domain of life (Eukarya, Bacteria, or Archaea) are used to maximize sensitivity for predicting different types of tRNA genes. The incorporation of the accelerated profile hidden Markov model (HMM) methods used in HMMER3 (25,26) and the constrained CM alignment algorithms (20,27) in Infernal provide multiple levels of filtering as part of sequence homology searches. In the default setting, tRNAscan-SE 2.0 adopts mid-level strictness by executing Infernal’s cmsearch with option “--mid” and a low score cutoff (10 bits) to replace the original first-pass filters in tRNAscan-SE 1.3. Then, for the second-pass high specificity scan, tRNAscan-SE 2.0 runs Infernal without the HMM filter on the first-pass candidates, including extra flanking sequences and a default score threshold of 20 bits. For users who need to obtain maximum search sensitivity and can accept a longer processing time, we also include the use of Infernal without HMM filter (--max option) as an alternative single-pass search option. It provides similar sensitivity as the COVE-only search mode in v1.3 but is significantly faster than the legacy method by at least 15 fold, depending on the genome size and number of repetitive elements (Supplementary Table S2). To further improve identification of slightly truncated tRNA genes, the new algorithm also makes use of the truncated hit detection feature in Infernal to annotate tRNA gene loci that may be truncated at either or both ends of the sequence. After initial tRNA gene prediction, the task of isotype classification is performed by comparing each anticodon prediction, identified by its position in the anticodon loop of the predicted tRNA secondary structure, to the scores of the full tRNA sequence against a suite of isotype-specific covariance models, a strategy similar to TFAM (28). Now, alongside the predicted anticodon, the highest scoring isotype-specific model can also be reported in the search output (--details option); any disagreement between the two functional prediction methods can be flagged by tRNAscan-SE to facilitate closer inspection by the user.

**Table 1.** Expanded phylogenetic representation of tRNA gene sets used as training for tRNA covariance model creation. The tRNA genes were grouped by source species’ domains for building domain-specific covariance models. For tRNAscan-SE 2.0, tRNA sequences were further sub-grouped into different isotypes for the generation of domain-specific, isotype-specific models.

| Domain | Models in tRNAscan-SE 1.3 |  | Models in tRNAscan-SE 2.0 |  |
| --- | --- | --- | --- | --- |
|  | No. of genera | No. of genomes | No. of genera | No. of genomes |
| Eukaryota | 88 | 115 | 110 | 155 |
| Bacteria | 23 | 33 | 647 | 4,016 |
| Archaea | 13 | 18 | 75 | 182 |
| Total | 124 | 166 | 838 | 4,285 |

**Figure 1.**
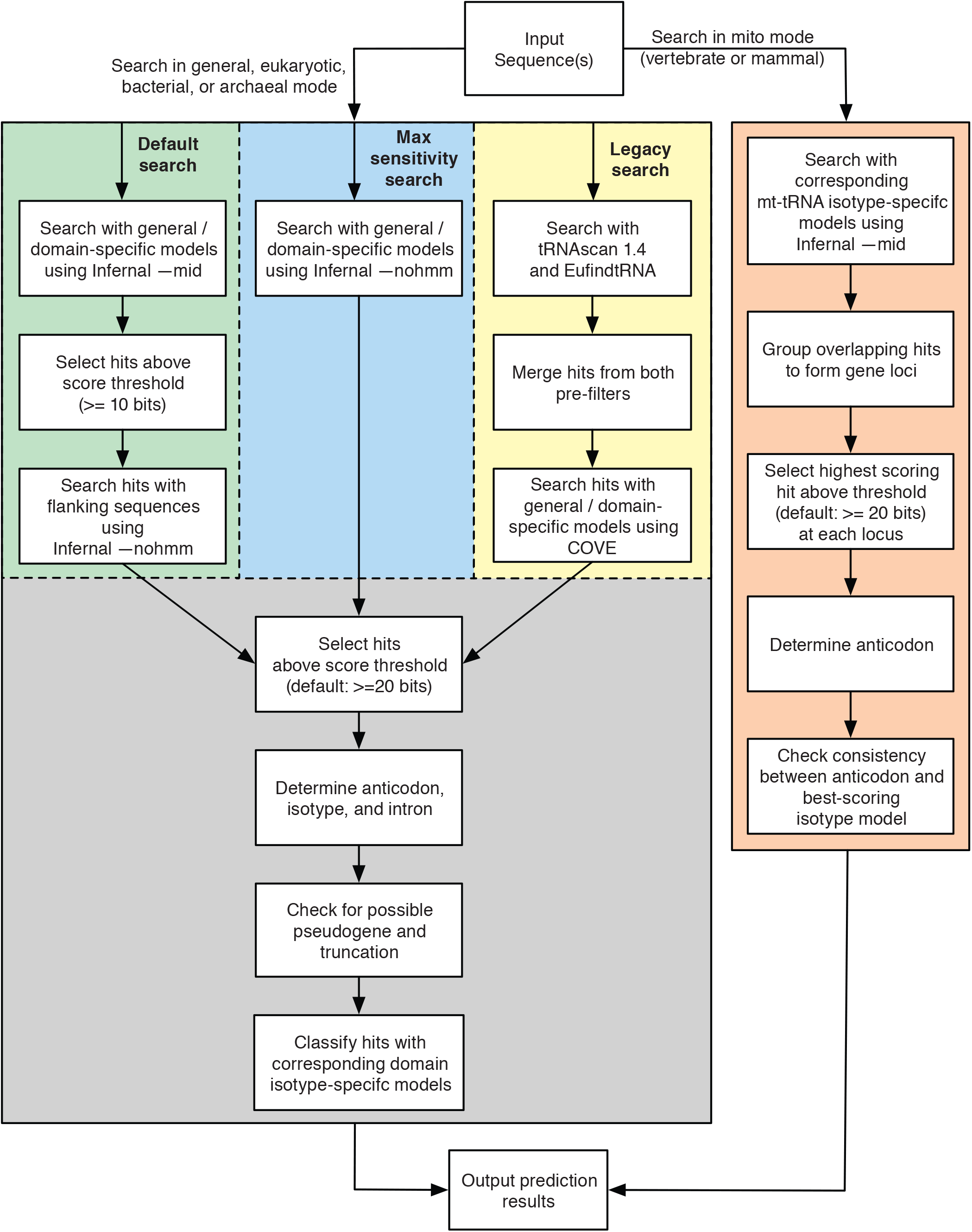
Schematic diagram of tRNAscan-SE 2.0 search algorithm. Three pathways were developed for cytosolic tRNA search modes with the addition of the mitochondrial tRNA search mode. The default method employs Infernal 1.1 (20) with newly built covariance models for similarity search while the legacy search remains the same as tRNAscan-SE 1.3 (1) for backward compatibility.

While tRNAscan-SE v1.3 included an organellar tRNA search option, it was not specifically trained on mitochondrial tRNAs, thus did not identify them with high accuracy, especially those that do not have a canonical cloverleaf secondary structure (29,30). In version 2.0, we have added in a mitochondrial tRNA search option (-M) that can use clade-and isotype-specific covariance models trained on mitochondrial tRNA (mt-tRNA) sequences. Currently, two versions of these mitochondrial search models are available: one set trained on mammalian mt-tRNAs, and another trained on a broader set of vertebrate mt-tRNAs. Additional mt-tRNA models are in development, particularly for species with highly degenerate mt-tRNA structures (e.g., missing one or more tRNA arms). The mitochondrial tRNA isotype indicated in prediction output corresponds to the highest scoring covariance model, while the anticodon is determined by analysis of the Infernal secondary structure output (middle three nucleotides found within the anticodon loop). This allows flagging of tRNA genes with inconsistent isotypes when comparing model-based versus anticodon-based isotype prediction. When scanning nuclear genomes, users can optionally include mitochondrial model scoring (--mt option) to identify and distinguish between cytosolic and nuclear-encoded mitochondrial tRNAs (31,32).

### False positive analysis and score threshold determination

The default score cutoff for tRNA predictions in tRNAscan-SE 1.3 using COVE (13) is 20 bits. To assess if this threshold could be applied to the new version with Infernal (20), we generated virtual genomes for *E. coli* K12, *Halobacterium sp*. NRC-1, *Saccharomyces cerevisiae* S288C, and *Homo sapiens* by using a 5^th^ order Markov chain to retain the base frequencies of actual genomes (Supplementary Table S3). These simulated sequences enable testing a broad diversity of G/C contents and genome sizes from evolutionarily distant phylogenetic clades. The tRNAscan-SE 2.0 default search mode and a score cutoff of 10 bits (“--score 10” option) was used to search the equivalent of 100 genomes of each of the four different species. The highest scoring hit in this negative control sequence set was 29.5 bits, and the vast majority of false positive hits (687 out of the total 704) had very low scores ranging between 10 and 20 bits (Supplementary Figure S2A). For reference, 100,700 tRNAs (100 genomes X 1007 total tRNA genes) were identified in the equivalent copies of actual genomes using a score cutoff of 20 bits or greater. Based on these results, we did not change the original 20-bit default score threshold used in prior versions of tRNAscan-SE. When scanning large eukaryotic genomes and loss of identification of some tRNA-derived pseudogenes is acceptable, a threshold of 30 bits (“--score 30” option) may be considered as a useful high-stringency cutoff.

Similarly, we applied this strategy for estimating false positive tRNA predictions in vertebrate mitochondria. 3,500 virtual mitochondrial genomes were generated with the same method as above for each of five genomes representing a range of G/C content (Supplementary Table S4). tRNAscan-SE 2.0 mitochondrial search mode for vertebrates (“-M vert” option) was used to search the virtual genomes. With the score cutoff set at 0 bit for this search, a total of 1,764 hits were identified with scores ranging between 0 and 16.5 bits (Supplementary Figure S2B). For reference, 539,000 tRNAs (3,500 genomes X 154 total mt-tRNA genes) were identified in the equivalent copies of actual genomes using 20 bits as the score cutoff. Thus, we kept a default cutoff of 20 bits for mitochondrial tRNA genes. We also evaluated the performance of ARWEN (8) and MiTFi (9) by scanning the same set of virtual genomes for comparison (Supplementary Table S4). ARWEN with -mtmam and -rp options were used for virtual genomes of *Homo sapiens* and *Hemiechinus auritus* while -gcvert and -rp were used for the rest. MiTFi was run in default mode.

### tRNAscan-SE 1.3 and 2.0 Search Comparisons

The genome assemblies used in this study were scanned with tRNAscan-SE v1.3 and 2.0 using the corresponding domain search modes (-E, -B or -A options). When running v1.3 on archaeal genomes, the --ncintron option was also included to enable noncanonical intron searches (this functionality is default in 2.0). Predicted tRNA genes in archaeal, bacterial, and fungal genomes were compared on the basis of similarity of genomic coordinates and tRNA identity (isotype and anticodon). Results were grouped into four categories: (1) consistent – the predicted gene has consistent identity and start and/or end positions differ by 10 nucleotides or less, (2) isotype mismatch – the predicted gene coordinates are the same but the isotype does not match, (3) novel – the gene is only predicted by v2.0 but not v1.3, (4) not detected – the gene is predicted by v1.3 but not v2.0. Within the “consistent” category (Table 2), only 0.69% archaeal, 6.8% bacterial, and 1.7% fungal tRNA predictions were found to have slightly different start and/or end positions. Model organisms (Table 3) were compared by predicted gene counts and program execution time. All prediction runs were conducted on Linux servers with identical configurations (dual Intel 10-core HT processors at 2.30GHz and 128 GB memory). tRNAscan-SE 2.0 utilized eight parallel threads for Infernal (20) searches (--thread 8 option) while v1.3 only allows a single thread.

**Table 2.** Comparison of tRNA gene predictions in archaeal, bacterial, and fungal genomes between tRNAscan-SE 1.3 and 2.0 show thousands of new tRNA identifications. tRNA gene counts and percentages given relative to v2.0 total predictions: “Consistent”, v2.0 and v1.3 gene predictions that are identical or nearly identical; “Isotype mismatch”, v2.0 and v1.3 isotype predictions that disagree for the same tRNA gene; “Novel”, v2.0 predictions that were not found by v1.3. “Not detected” includes v1.3 predictions not found by v2.0 (percentages are relative to v1.3 total predictions).

| Clade | # of genomes | tRNAscan-SE 1.3 predictions | tRNAscan-SE 2.0 predictions | Consistent | Isotype mismatch | Novel | Not detected |
| --- | --- | --- | --- | --- | --- | --- | --- |
| Archaea | 217 | 10,302 | 10,354 | 10,205<br>(98.5%) | 79<br>(0.8%) | 70<br>(0.7%) | 29<br>(0.28%) |
| Bacteria | 4,036 | 240,352 | 246,226 | 239,612<br>(97.3%) | 447<br>(0.2%) | 6,165<br>(2.5%) | 293<br>(0.12%) |
| Fungi | 432 | 81,620 | 80,824 | 74,997<br>(92.8%) | 3,398<br>(4.2%) | 2,426<br>(3.0%) | 3,225<br>(3.9%) |

**Table 3.** Comparison of tRNAscan-SE predictions counts and search performance for selected model species. (A) tRNA prediction counts for v1.3, and relative change for v2.0, show no difference or small gains in bacteria and archaea. In eukaryotes with large numbers of pseudogenes, the differences between v1.3 and v2.0 are much reduced after excluding pseudogenes. (B) Search times of genomes using v1.3 and v2.0.

A.
| Genome | Domain | Genome size (million bp) | tRNAscan-SE 1.3 / 2.0 |  |
| --- | --- | --- | --- | --- |
|  |  |  | Total tRNA predictions | tRNAs excluding pseudogenes |
| *Bacillus subtilis* 168 | Bacteria | 4.2 | 86/0 | 86/0 |
| *Clostridioides difficile* 630 | Bacteria | 4.3 | 87/+2 | 87/+1 |
| *Escherichia coli* K-12 | Bacteria | 4.6 | 88/+1 | 87/+1 |
| *Mycobacterium tuberculosis* H37Rv | Bacteria | 4.4 | 45/0 | 45/0 |
| *Pseudomonas aeruginosa* PAO1 | Bacteria | 6.3 | 64/0 | 63/0 |
| *Halobacterium* sp. NRC-1 | Archaea | 2.0 | 47/0 | 47/0 |
| *Methanocaldococcus jannaschii* | Archaea | 1.7 | 37/0 | 37/0 |
| *Pyrobaculum aerophilum* | Archaea | 2.2 | 45/+2 | 45/+2 |
| *Pyrobaculum calidifontis* | Archaea | 2.0 | 43/+3 | 43/+3 |
| *Pyrococcus furiosus* DSM 3638 | Archaea | 1.9 | 46/0 | 46/0 |
| *Sulfolobus acidocaldarius* DSM 639 | Archaea | 2.2 | 49/+1 | 49/+1 |
| *Saccharomyces cerevisiae* | Eukaryota | 12.2 | 275/0 | 275/0 |
| *Schizosaccharomyces pombe* 972h- | Eukaryota | 12.6 | 171/0 | 171/0 |
| *Homo sapiens* | Eukaryota | 3,101 | 625/-29 | 515/-4 |
| *Mus musculus* | Eukaryota | 2,731 | 26,287/+14,625 | 3,296/+1,201 |
| *Caenorhabditis elegans* | Eukaryota | 100.3 | 820/-99 | 609/+9 |
| *Drosophila melanogaster* | Eukaryota | 143.7 | 295/0 | 291/+1 |
| *Arabidopsis thaliana* | Eukaryota | 119.7 | 639/+3 | 631/+3 |
| *Zea mays* | Eukaryota | 2,134 | 2,296/-213 | 1,464/-76 |

B.
| Genome | Domain | tRNAscan-SE 1.3 | tRNAscan-SE 2.0 |
| --- | --- | --- | --- |
| *Bacillus subtilis* 168 | Bacteria | 27s | 37s |
| *Clostridioides difficile* 630 | Bacteria | 19s | 37s |
| *Escherichia coli* K-12 | Bacteria | 43s | 48s |
| *Mycobacterium tuberculosis* H37Rv | Bacteria | 40s | 23s |
| *Pseudomonas aeruginosa* PAO1 | Bacteria | 56s | 33s |
| *Halobacterium* sp. NRC-1 | Archaea | 1m 35s | 51s |
| *Methanocaldococcus jannaschii* | Archaea | 29s | 42s |
| *Pyrobaculum aerophilum* | Archaea | 1m 10s | 1m 19s |
| *Pyrobaculum calidifontis* | Archaea | 1m 43s | 2m 9s |
| *Pyrococcus furiosus* DSM 3638 | Archaea | 33s | 57s |
| *Sulfolobus acidocaldarius* DSM 639 | Archaea | 47s | 1m 1s |
| *Saccharomyces cerevisiae* | Eukaryota | 52s | 2m 20s |
| *Schizosaccharomyces pombe* 972h- | Eukaryota | 36s | 1m 24s |
| *Homo sapiens* | Eukaryota | 1h 37m | 49m 37s |
| *Mus musculus* | Eukaryota | 7h 46m | 4h 33m |
| *Caenorhabditis elegans* | Eukaryota | 5m 9s | 5m 39s |
| *Drosophila melanogaster* | Eukaryota | 3m 54s | 6m 12s |
| *Arabidopsis thaliana* | Eukaryota | 4m 45s | 5m 31s |
| *Zea mays* | Eukaryota | 1h 5m | 46m 39s |

### Development of domain-specific covariance models

We assembled three sets of domain-specific genomic tRNA sequences from a total of over 4,000 genomes using existing tRNAscan-SE 1.3 predictions in GtRNAdb Release 15 (4) plus additional predictions (also from tRNAscan-SE 1.3 for species not yet represented in GtRNAdb), representing a broad diversity of eukaryotes, bacteria, and archaea (Table 1, Supplementary Figure S1). Before using these tRNA sequences as training sets for building domain-specific covariance models, multiple filtering steps were used to maximize quality. To avoid the inclusion of common tRNA-derived repetitive elements that exist in many eukaryotic genomes, especially those in mammals (33–35), we first selected only the eukaryotic tRNAs with a COVE score greater than 50 bits, a threshold reflecting more conserved, canonical tRNA features. We then selected only the top 50 scoring tRNAs for each isotype per organism to avoid overrepresentation of high-scoring tRNA-derived repetitive elements which are abundant in some species (for example, elephant shark has over 9,500 tRNA^Ala^ scoring over 50 bits). For the bacterial tRNA training set, all genes having potential self-splicing introns were excluded to eliminate large alignment gaps (i.e., mostly over 200 nt) which can hinder efficient model creation. Similarly for archaea, pre-processing of sequence training sets was necessary. Some species within the phyla Crenarchaeota and Thaumarchaeota contain tRNAs that are known to have multiple noncanonical introns (5,36,37). Atypical tRNAs such as *trans*-spliced tRNAs and circularized permuted tRNAs have also previously been discovered in Crenarchaeota and Nanoarchaeota (12,38,39). To accommodate these special archaeal features without sacrificing performance, both mature tRNA sequences (without introns) and selected atypical genes with multiple introns at different locations were included in the archaeal tRNA training sets. As a last step, anticodons of tRNA sequences in all training sets were replaced with NNN and aligned to the corresponding original domain-specific tRNA covariance models using Infernal (20). The resulting alignments were then used to generate the new set of domain-specific tRNA covariance models with Infernal.

### Development of isotype-specific covariance models

The isotype-specific covariance models for the three phylogenetic domains were built iteratively through two rounds of training to capture sequence features identifying each tRNA isotype (Supplementary Figure S1). We found this two-step process necessary to avoid polluting isotype-specific alignments with tRNAs derived from other isotypes but “mislabelled” by the prior tRNAscan-SE due to mutations in anticodons (relatively common in high-scoring pseudogenes). In the first round, the tRNA genes used for building the domain-specific covariance models were divided into groups according to the tRNA isotypes determined by the anticodon sequence. For archaeal models, tRNA sequences of genes with noncanonical introns were “pre-spliced” to only include their mature sequences. The sequences of each isotype group were then aligned to the tRNA covariance model of the corresponding domain using Infernal (20). “Intermediate” covariance models for each isotype were built using the resulting alignments. In a second round of model-building, the original training set from each domain was scored against the new intermediate covariance models. The sequences were then grouped according to isotype, this time primarily based on which isotype-specific covariance model yielded the highest score for each sequence (regardless of the anticodon sequence), followed by manual inspection and adjustments if needed to ensure correctness. These revised isotype-classified sequence groups were re-aligned to the corresponding intermediate models to build the final isotype-specific covariance models.

### Covariance models for methionine tRNAs and isoleucine tRNAs decoding AUA

In eukaryotes and archaea, initiator methionine tRNA (tRNA^iMet^) and elongator methionine tRNA (tRNA^Met^) have distinct sequence features and functions but contain the same anticodon, CAU. Similarly, N-formylmethionine tRNA (tRNA^fMet^) in bacteria also contains the same CAU codon as the structurally distinct elongator methionine tRNA. Further adding to anticodon-function ambiguity, tRNA^Ile2^, which decodes the isoleucine AUA codon, is also encoded by tRNA genes containing a CAU anticodon. These special tRNA^Ile2^ are post-transcriptionally modified at their wobble bases (lysidine in bacteria and agmatidine in archaea), effectively giving them isoleucine-specific UAU anticodons. The strategy of prior versions of tRNAscan-SE to identify tRNAs based only on their anticodon failed to separate these three functionally distinct tRNAs.

In order to develop accurate covariance models that represent these structurally and functionally different tRNAs, we applied the above two-round training method with carefully selected training sets. For eukaryotes, the sequences of tRNA^iMet^ and tRNA^Met^ were selected from the original 1,415 tRNAs used for training tRNAscan-SE 1.3. For bacteria, we collected the sequences for tRNA^fMet^, tRNA^Met^, and tRNA^Ile2^ from 234 genomes where these tRNAs were classified (40). For archaea, we curated the sequences based on known identity elements of these three different tRNAs (41). These sequences were aligned to the corresponding domain-specific covariance models for generating the first-round intermediate covariance models followed by the second-round training step, producing the final covariance models.

### Selenocysteine tRNA modelling

Selenocysteine tRNAs (tRNA^SeC^) have secondary structures and loop sequence lengths that differ from other tRNA isotypes, and are a component of the protein translation system of species that employ selenoproteins (42). While eukaryotic and archaeal tRNA^SeC^ have a 9-bp acceptor stem and a 4-bp T-arm (9/4 fold) (43–46), bacteria have an 8-bp acceptor stem and a 5-bp T-arm (8/5 fold) (47,48). To build covariance models for these special cases, we curated the sequences of tRNA^SeC^ from genomes where selenoproteins were previously identified and removed those that do not retain published canonical features: the special secondary structure of tRNA^SeC^, a UCA anticodon, and G at position 73. The collected sequences (65 eukaryotic (49–60), 61 bacterial (61), and 13 archaeal (46,62–65) were aligned to the original tRNA^SeC^ covariance models from tRNAscan-SE 1.3, with manual inspection and adjustments. The resulting alignments were used to build the domain-specific tRNA^SeC^ covariance models.

### Identification of noncanonical introns in archaeal tRNAs

All tRNA candidates predicted with the archaeal search model are analyzed for noncanonical introns (Supplementary Figure S3). Two covariance models that include the bulge-helix-bulge (BHB) secondary structure were built with manually curated noncanonical tRNA introns from (1) Crenarchaeota and Euryarchaeota, and (2) Thaumarchaeota, in order to capture the different consensus intron sequences found in tRNAs from these phylogenetic groups. To improve the identification of introns located near the 5’ and/or 3’ ends, tRNA candidates are extended with 60 nucleotides of flanking sequences and scanned with the BHB covariance models. Predicted noncanonical introns are confirmed when the score of the predicted mature tRNA with detected intron(s) removed is higher than the unspliced form. Some tRNAs contain two or three introns, with introns located in such close proximity that one intron must be removed before a second BHB motif can be formed and detected in the second intron. Thus, multiple iterations of the intron search are carried out until the tRNA prediction score is maximized, or the length of the predicted mature tRNA is less than 70 nucleotides (the typical minimum length of archaeal tRNAs). Final reported scores in tRNAscan-SE 2.0 outputs are based on the predicted mature tRNAs.

### Covariance models for detecting mitochondrial tRNAs in vertebrates

Mitochondrial tRNAs (mt-tRNAs) in 1,085 vertebrate species were obtained from mitotRNAdb (17) as the training set for building covariance models. In the mitochondria of vertebrates, there is typically one tRNA for each isotype, except mt-tRNA^Leu^ and mt-tRNA^Ser^ which each have two tRNAs with different anticodons (66). To achieve high specificity for each mt-tRNA type, we grouped the sequences by isotypes and anticodons resulting in 22 sets of mt-tRNAs for generating individual covariance models. To further increase prediction accuracy for mammalian mt-tRNAs, we built a second set of covariance models that only utilized mt-tRNAs from 282 mammals as the training set. Both sets of covariance models were built with two rounds of training. First, each subset of isotype-specific mt-tRNA sequences, excluding mt-tRNA^Ser(GCU)^, was aligned to the general covariance model from the original tRNAscan-SE (1) using Infernal (20). The general model was chosen because it was used in the organellar search mode of tRNAscan-SE v1.3 and has the greatest diversity of tRNA training sequences from all three domains of life. To accommodate the atypical secondary structure of mt-tRNA^Ser(GCU)^, a custom covariance model without the D-arm was built and used for aligning mt-tRNA^Ser(GCU)^ genes. The resulting alignments were used to build the first set of 22 isotype-specific covariance models for vertebrates and mammals. For the second round of training, the isotype-specific mt-tRNA sequences were aligned to the corresponding isotype-specific covariance models generated in the first round. These alignments were then used to build the final sets of mt-tRNA covariance models.

### Prediction evaluation for mitochondrial tRNAs in vertebrates

NCBI RefSeq (24) mt-tRNA gene annotations from 3,345 vertebrate mitochondrial genomes that include 739 mammals (genome size range: 9118 to 28757 bytes; median size: 16616 bytes; average size: 16802 bytes) (Supplementary File S7) were obtained to compare with the tRNAscan-SE 2.0 predictions using the vertebrate mitochondrial search mode in the corresponding genomes; the mammalian-specific mitochondrial models showed similar performance, and therefore all subsequent analyses used the more diverse vertebrate mt-tRNA models for easier comparison. For testing purposes, the score cutoff of tRNAscan-SE was set to 0 bits to better identify potential “false negatives” scoring just below the standard 20 bit cutoff (Supplementary Figure S4 and Supplementary File S8). Comparison results were grouped into five categories: (1) consistent – the predicted gene overlaps within 15 nucleotides of the start and end positions and also has a consistent isotype as annotated in RefSeq, (2) isotype mismatch – the predicted gene is located in the same genomic locus as RefSeq annotation but the isotype does not match, (3) position mismatch – the predicted gene has identical isotype as RefSeq annotation but strandedness is different or the predicted gene boundaries are beyond +/-15 nucleotides (although still overlapping with RefSeq annotation), (4) novel – the predicted gene does not have a corresponding RefSeq annotation in the same genomic locus, (5) not detected – RefSeq-annotated gene not detected by tRNAscan-SE. Predictions from tRNAscan-SE 2.0 that were not classified as matches were inspected to assess the accuracy of the program and/or the existing annotation.

For algorithm performance comparison, mt-tRNA predictions in the same set of vertebrate mitochondrial genomes were generated using ARWEN (8) and MiTFi (9). ARWEN was run with recommended options of –mtmat for mammalian mitochondrial genomes and –gcvert for non-mammalian vertebrate mitochondrial genomes; MiTFi was run with default mode. Predictions from each algorithm were also compared against NCBI RefSeq (24) gene annotations and comparison results were grouped into categories as described above. These results were compared with predictions of tRNAscan-SE 2.0 using the default score cutoff (20 bits).

### Custom search configurations

To increase the flexibility of tRNAscan-SE 2.0, we include a configuration file that accompanies the software. This file contains default parameters such as score thresholds of different search modes, file locations of covariance models, and legacy search mode settings. Advanced users can make changes to settings as appropriate for their research needs. To further extend its capability, an extra search option was implemented in tRNAscan-SE 2.0 that allows researchers to use alternate covariance models specified in the configuration file for tRNA searching. If multiple alternate covariance models are included, the top scoring one at each overlapping locus will be reported. This new feature enables the use of custom-built covariance models that may better detect tRNAs with atypical unique features.

### Post-scan filtering of potential tRNA-derived repetitive elements and classification of high confidence tRNAs

EukHighConfidenceFilter in tRNAscan-SE 2.0 is a post-scan filtering program for better distinguishing tRNA-derived repetitive elements in large eukaryotic genomes (metazoans and plants) from “real” tRNAs that function in protein translation. Three filtering stages are involved in the classification (Supplementary Figure S5). First, tRNA predictions that are labeled as possible pseudogenes are excluded from the high confidence set; these criteria were established by the prior version of tRNAscan-SE (overall score below 55 bits *and* one of two conditions: primary sequence score below 10 bits or secondary structure score below 5 bits). Second, predictions with *any* of the following attributes are removed from the high confidence set: isotype-specific model score below 70 bits, overall score below 50 bits, or secondary structure score below 10 bits. Finally, if there are more than 40 predicted hits remaining for any given anticodon, a dynamic score threshold is used, starting at 71 bits, rising one bit and filtering lower-scoring hits, iteratively, until the number of predictions for that anticodon is no longer over 40 *or* when the score threshold reaches 95 bits. The score thresholds in stages two and three were empirically determined by comparing score distributions of predictions among eukaryotic genomes with and without large numbers of tRNA-derived repetitive elements (Supplementary Figure S6). The remaining tRNA predictions are included in the high confidence set if (1) they have a consistent isotype prediction (inferred from anticodon versus the highest scoring isotype-specific model) *and* (2) they have an “expected” anticodon based on known decoding strategies in eukaryotes where 15 anticodons are not used (67). There is no corresponding filter for bacterial or archaeal tRNAs, as they have not been found to contain large families of tRNA-derived repetitive elements.

## RESULTS

### Improvements in performance of tRNAscan-SE 2.0

With thousands of genomes that have become available since the first tRNAscan-SE was released, there is an opportunity to improve the diversity of sequences used to train its search models, ensuring high sensitivity for an ever-broader group of sequenced organisms. Genome sequencing has far eclipsed direct tRNA sequencing for many years, thus the vast majority of observed tRNA sequences are no longer based on tRNA transcripts. Instead, they are almost always computationally predicted from genomes, most likely by an earlier version of tRNAscan-SE. As such, the first step in assessing the algorithm performance of tRNAscan-SE 2.0 (Figure 1) is to compare it to the existing gold standard, tRNAscan-SE 1.3. One of the most substantial advancements in the new version is incorporation of Infernal 1.1 (20) (see Methods), which makes use of multi-core central processing units (CPUs) that enable parallelization to achieve accelerated searches (26). Integration of Infernal also enables elimination of the “first-pass” scanning programs (Figure 1, “Legacy search”) required for speed in tRNAscan-SE 1.3. This algorithmic streamlining provided new flexibility to develop a broader suite of specialized tRNA covariance models. New and updated models now include tRNA training data from over 4000 genomes across the three domains of life (Table 1). As a result, tRNAscan-SE 2.0 shows improvement in overall detection sensitivity, increased accuracy in detection of particular classes of tRNAs, and a new way to better predict the tRNA isotype, all at little to no cost in execution time relative to the prior version.

Comparing detection results between tRNAscan-SE 1.3 and 2.0 across a wide swath of high-quality complete prokaryotic genomes, we found 10,205 archaeal (98.5%) and 239,672 bacterial (97.3%) predictions that were identical or substantially overlapping (Table 2), leaving a relatively small proportion with differences to be examined more closely. We noted these changes were among relatively weak tRNA predictions (scores of <50 bits) for which normal tRNA processing and function is uncertain. All differences were broken down into three classes: changes in tRNA isotype, predictions only detected using v1.3, and new predictions found only using v2.0.

Among the 178 archaeal tRNAs showing differences (Table 2), the majority of these were low-scoring (< 50 bits), including all 29 tRNAs no longer reported by v2.0 (Figure 2A). Manual inspection showed that these low scores were often caused by truncated genes (either on 5’ or 3’ end) or noncanonical cloverleaf secondary structures (Supplementary File S1), suggesting that they may have little or no role in protein translation. Conversely, improvements in v2.0 for detecting tRNAs with noncanonical introns (see Methods) enables identification of more than three dozen high-scoring archaeal tRNAs that were previously missed, mostly in species from the phyla Crenarchaeota and Thaumarchaeota (5,36,37,68) (Supplementary File S1). Thus, v2.0 shows clear improvement in sensitivity for the most difficult tRNA targets previously overlooked. As more archaeal species with non-canonical introns are isolated and studied, this improvement will become even more essential for accurate, complete archaeal genome analyses.

**Figure 2.**
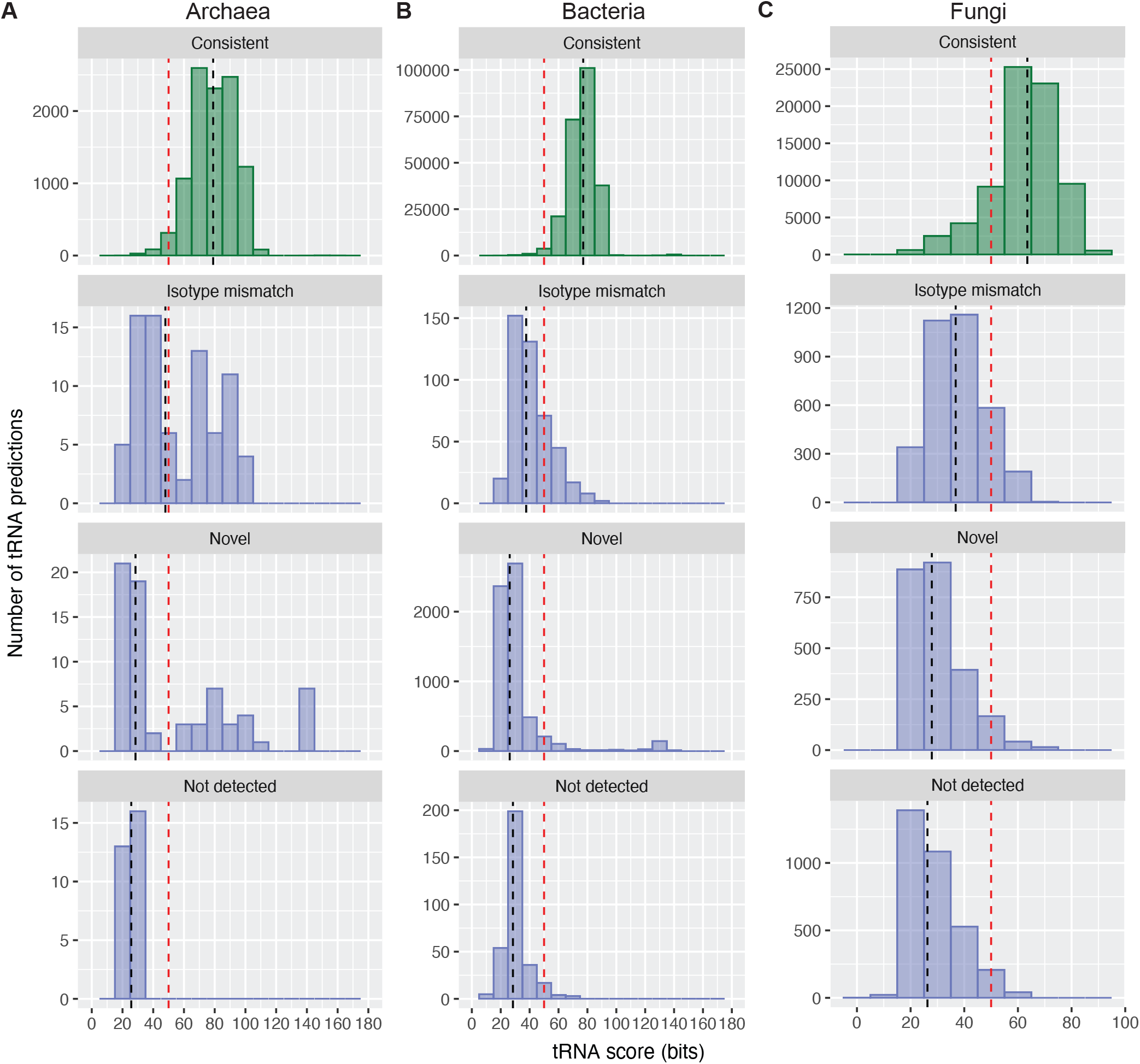
Score distributions of tRNAs comparing tRNAscan-SE 2.0 to 1.3 predictions show substantial agreement, with most differences among low-scoring (<50 bit) tRNAs. (A) 10,354 tRNA predictions in 217 archaeal genomes, (B) 246,271 predictions in 4,036 bacterial genomes, and (C) 80,824 predictions in 432 fungal genomes were detected using tRNAscan-SE 2.0 with default search modes for each respective domain. Prediction results were grouped into four categories: (1) “Consistent” between versions, (2) “Isotype Mismatch” where v2.0 and v1.3 predictions have different isotype classifications, (3) “Novel” for predictions new in v2.0, and (4) “Not detected” for prior 1.3 predictions no longer detected by v2.0. Each histogram represents the prediction scores binned by 10 bits in the corresponding comparison category. Vertical black dashed lines represent the median score in the category, and vertical red dashed lines demarcate the 50 bit “low score” threshold.

Within bacterial genomes, there were almost 7,000 tRNA genes with detection differences between v1.3 and v2.0 (Table 2), but 6,165 of these were new predictions found only by v2.0 (Table 2). A majority of these detection differences occurred in low-scoring tRNAs (Figure 2B) — for example, the median score for predictions that matched between versions was 77.1 bits, whereas the medians were 28.4 bits for those not detected by v2.0, and 26.2 bits for novel v2.0 predictions. Similar to those in archaea, we estimated that these low-scoring tRNA predictions are not transcribed or normally processed, thus would have minimal impact on protein translation. Of all the newly identified bacterial tRNAs, selenocysteine tRNAs (tRNA^SeC^) stand out, as these tRNA genes are now detected by v2.0 as high-scoring hits in 273 bacterial genomes due to the greatly improved tRNA^SeC^ covariance models (see Methods). Of the 293 bacterial tRNA predictions detected by v1.3 but not v2.0, all were predicted by v1.3 to be pseudogenes, had atypical secondary structures and/or scored less than 50 bits (Supplementary File S2). Our analyses of more than 240,000 bacterial tRNAs demonstrate a clear net gain in bacterial tRNA prediction for v2.0.

While we observed mostly identical but slightly improved detection in the new version of tRNAscan-SE for bacteria and archaea, there were many more differences in eukaryotes due to dramatically increased genome sizes, incompletely assembled chromosomes, and increased frequency of tRNA-derived repetitive elements. For this reason, we utilized fungi for benchmarking, the eukaryotic subdomain that has the largest number of complete, relatively compact genomes which harbor smaller numbers of tRNA-derived repetitive elements in contrast to many vertebrate genomes (33,69). As described for bacteria and archaea, the vast majority of inconsistencies between v1.3 and v2.0 were from sub-50 bit tRNA gene predictions (Table 2, Figure 2C, Supplementary File S3). The median score for fungal predictions that matched between versions was 63.5 bits, whereas the medians were 26.3 bits for those not detected by v2.0, and 27.9 bits for novel v2.0 predictions. For tRNAs showing differences in isotype prediction, the median score was 36.8 bits; noncanonical cloverleaf secondary structures in these tRNAs, found in a small subset of fungal genomes with tRNA-derived repetitive elements (69–71), accounted for nearly all these differences in isotype prediction. Overall, the approximately 93% consistency between v1.3 and 2.0 predictions for 432 fungal genomes, with nearly all conflicts occuring in low-scoring hits, shows remarkably similar performance for typical tRNAs (Figure 2C).

### Effect of New Algorithm on tRNA Detection in Model Genomes

Focusing more specifically on model species (Table 3), we investigated the sequence basis for every difference in predictions between versions. For *Escherichia coli* K-12, we noted an extra predicted tRNA gene was identified by tRNAscan-SE 2.0, although it is very low-scoring (29.6 bits) with an undetermined anticodon and isotype (Supplementary File S2). Upon inspection, we found that it is part of gene b2621, a transfer-messenger RNA (tmRNA) that is known to have both tRNA-like and mRNA-like properties and rescues stalled ribosomes (72,73). In other well-studied bacterial species, multiple tRNA genes in *Bacillus subtilis* 168 (2 genes) and *Mycobacterium tuberculosis* H37Rv (15 genes) were predicted to be 3 nucleotides shorter using the new version. Comparison of the sequence alignments revealed that tRNAscan-SE 1.3 incorrectly included additional trailer sequences such as TCA and CTA as the 3′ CCA tail, indicating an improvement of the newly built covariance models for identifying the 3’ ends of bacterial tRNAs in v2.0. In addition, the better-trained model for bacterial tRNA^SeC^ enables correct prediction of this tRNA’s gene boundaries in *Pseudomonas aeruginosa* PAO1 and avoids v1.3’s omission of the gene in *Clostridioides difficile* 630 (heterotypic synonym: *Peptoclostridium difficile* 630).

For model archaea such as *Pyrococcus furiosus, Sulfolobus acidocaldarius, Halobacterium sp*. NRC-1, *and Methanocaldococcus jannaschii*, the predicted genes were identical or had a slight improvement in sensitivity for intron-containing genes (Table 3A, Supplementary File S1). Similarly, there were no changes in the number of predicted genes for the model single-celled fungi *Saccharomyces cerevisiae* S288C (budding yeast) and *Schizosaccharomyces pombe* 972h-. In *Drosophila melanogaster*, tRNAscan-SE 2.0 correctly detects one more tRNA gene (tRNA-Tyr-GTA-1-9) that was originally missed by v1.3 due to its large 113-nt intron, while a likely pseudogene with a very low score (28.6 bits) in the genome is dropped by v2.0 (Supplementary File S3). By comparison, *Caenorhabditis elegans* showed more differences than *Drosophila*, yet 95% (685 out of 721) of the predictions by v2.0 were consistent with v1.3. Among the 14 predictions that had different isotype/anticodon, eleven of them were marked as pseudogenes and the others had atypical predicted secondary structures or sequence features. The remaining *C. elegans* differences were again among predicted pseudogenes and other sub-50 bit predictions, with one notable exception. Similar to *Drosophila*, one of the novel high-scoring genes found exclusively by v2.0 (tRNA-Ala-AGC-7-1) was predicted to have a very large intron (323 nt). Thus, v2.0 is estimated to have the same or slightly better sensitivity for well-studied model species.

### Human tRNA Detection

Examination of the changes in human tRNA predictions further illustrates subtle differences seen in other vertebrate analyses, mostly among borderline predictions. Out of 625 prior tRNA predictions in the human genome, tRNAscan-SE 2.0 detects 56 more candidates but does not report 65 predicted by v1.3. Within these non-intersecting sets, two-thirds are classified as possible pseudogenes by tRNAscan-SE (see Methods), with the highest score among predictions found only by v2.0 at 47.6 bits, or only found by v1.3 at 34.44 bits. To gather experimental evidence for transcription and/or processing of these low-scoring predictions, we examined published small RNA sequencing data from ARM-Seq (74), DM-tRNA-seq (75,76), and the DASHR database (77). Based on tRNA read abundance from these studies (Supplementary Table S5), we found that only two of the predictions detected only by v1.3 are above background (>5 reads per million, RPM; Supplementary Table S5A), and these reads only aligned to the 3’ end of the tRNA gene (Supplementary Figure S7). Among the highest scoring new v2.0 predictions examined, nine showed expression above background in at least one sample (Supplementary Table S5B). Most genes showing some expression have sequencing reads aligned to just the 5’ or 3’ end of the gene body, suggesting possible roles as tRNA-derived small RNAs. Two predictions show a different transcript pattern, with RNA-seq reads aligned to the anticipated full length pre-tRNA transcript. One of them is a low-scoring (22.8 bits) tRNA^Tyr(GTA)^ gene with a 12-nt intron and low abundance level across the entire precursor tRNA region (Supplementary Figure S8A) (74). Based on the UCSC Genome Browser 100-vertebrate conservation track (22), this tRNA-like feature aligns to similarly low-scoring versions in primates but also high-scoring canonical tRNA^Tyr(GTA)^ genes in non-primate mammals. Recent mutations in the primate lineage suggest it has become a tRNA-derived element that is still transcribed but not processed into a functional tRNA (Supplementary Figure S8B). Overall, our survey of human tRNA sequencing data revealed a small fraction of these “new” predictions (all <50 bits) that have evidence of transcription, yet apparently not as stable full-length tRNAs directly involved in ribosomal codon translation. Throughout the comparisons between the two versions of the program, this was a recurring theme: very few changes for high confidence tRNAs with evidence of expected function, versus numerous changes in “edge cases” in the twilight region (∼20-50 bits) where tRNA-derived gene remnants retain some but not all the sequence features necessary for transcription, processing to maturity, and deployment for ribosomal translation.

### False Positive and Speed Comparison

To assess false positive rate, we used tRNAscan-SE 2.0 to search against the equivalent of 100 synthetic genomes with preserved hexamer frequencies for each of four different model species (representing varied G/C content) including *E. coli* K12, *Halobacterium sp*. NRC-1, *Saccharomyces cerevisiae* S288C, and *Homo sapiens* (see Methods). No false positives were detected above 20 bits in the three sets of virtual microbial genomes, and only 17 hits were found above the default 20-bit cutoff in the virtual human genomes (Supplementary Table S3 and Supplementary File S4). For context, 100,700 tRNAs were identified in the equivalent number of copies of the actual model genomes. The highest hit scored 29.5 bits (Supplementary Figure S2A), which is substantially lower than the median score for typical tRNA genes (Figure 2), and all 17 would be classified as pseudogenes. The original version of tRNAscan-SE previously reported a false positive rate of < 0.00007 per million bp (1). Thus, with a false positive rate of 0.00005 per million bp, v2.0 is estimated to have an equal or slightly better selectivity than v1.3.

We compared the relative execution speed of both versions of the program on model genomes (Table 3B). tRNAscan-SE 2.0 is slightly slower than v1.3 for organisms with small genome size (e.g., *E. coli* and *Saccharomyces cerevisiae*) due to the 20-fold isotype-specific covariance model analyses, a key new feature giving better functional classification (described below). However, the run time for large eukaryotic genomes which have a lower density of tRNA genes is much faster. For example, searching the human and mouse genomes with v2.0 with eight computing cores (see Methods) reduces the processing time by over 40%. Just as with the prior version, large genomes that have thousands of tRNA-derived repetitive elements are slower (e.g., mouse, 4h 33min for v2.0) and those with few of these elements are much faster (e.g., human, 49min for v2.0).

### Effective discrimination between initiator methionine, elongator methionine, and isoleucine 2 tRNAs

When tRNAscan-SE was originally designed, only a limited number of tRNA genes were identified and available in the 1993 Sprinzl database (16). The number and diversity of these sequences was not sufficient to train robust isotype-specific covariance models. However, with the multitude of complete genomes currently available, specialized models for individual functional classes of tRNAs can now be created. Each of the aminoacyl tRNA synthetase enzymes use sequence-and structure-based positive and negative determinants to ascertain tRNA identity; these features have been characterized in a number of model species (78). For simplicity, the original version of tRNAscan-SE used the anticodon exclusively to predict isotype because anticodon-isotype pairings are highly conserved and the anticodon is usually easy to determine in predicted tRNA genes. However, this method does not provide the capability to distinguish between different tRNA types which contain the same anticodon. The most ubiquitous example of this is the group of tRNA genes with the anticodon CAT in their genomic sequences — these include initiator methionine/N-formylmethionine tRNAs (tRNA^iMet^ in archaea/eukaryotes or tRNA^fMet^ in bacteria), elongator methionine tRNAs (tRNA^Met^ in all species), and isoleucine tRNAs decoding the AUA codon (tRNA^Ile2^ in bacteria and archaea).

By employing specialized models for these sub-groups of tRNAs in each domain of life, tRNAscan-SE 2.0 is now able to distinguish them (Figure 3). When comparing the scores of the different isotype-specific covariance models for tRNAs across all three domains of life (see Methods), we found that the tRNA^iMet/fMet^, tRNA^Met^, and tRNA^Ile2^ form distinct clusters (Figure 3), with the tRNA^iMet/fMet^ group the best-separated from the other two in bacteria and archaea. This reflects tRNA^iMet/fMet^ translation-initiation features that are distinct from tRNA^Met^ and tRNA^Ile2^ (41). To assess overall sensitivity of the new v2.0 isotype models, we checked the number of identified tRNAs in each studied genome (excluding incomplete fungal genomes), expecting at least one of each CAT anticodon subtype. Nearly all genomes contained the expected tRNAs, leaving 3.5% (6 of 173) eukaryotes, 1.8% (4 of 217) archaea, and 3.5% (143 of 4036) bacteria that were missing at least one gene of each subtype. Of the eukaryotic genomes with “missing” CAT gene predictions (four human pathogenic fungal species from the genus *Encephalitozoon*, and two biotrophic pathogens of *Zea mays*), all appear to have their tRNA^iMet^ mis-labeled as tRNA^Met^ because the iMet covariance model scores second best, just below the elongator Met model (Supplementary File S5). This small fraction of tRNA sequences from select fungal pathogens contain other atypical tRNA features when compared to the eukaryotic consensus, but were manually identified based on special features of eukaryotic iMet tRNAs that are mostly absent in elongator Met tRNAs such as A1:U72 base pair at the end of the acceptor stem (79).

**Figure 3.**
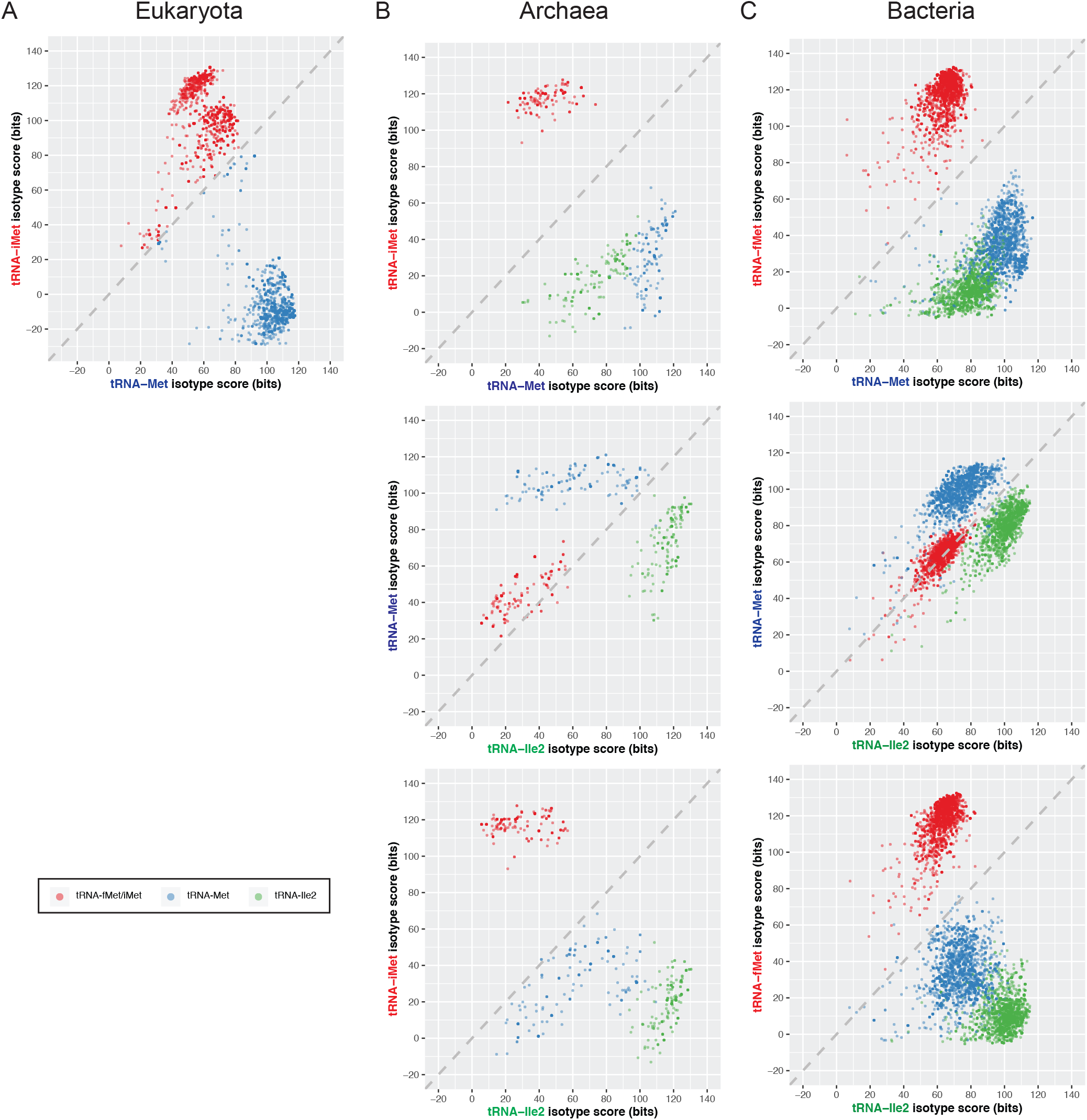
Direct comparison of isotype-specific covariance model scores provides an effective classifier of tRNAs with anticodon CAU. Dots represent individual tRNAs with the highest scores from the initiator methionine/N-formylmethionine model (tRNA^iMet/fMet^, red), elongator methionine model (tRNA^Met^, blue), or isoleucine-decoding CAU model (tRNA^Ile2^, green). Each tRNA was scanned with the isotype-specific covariance models of the corresponding genome’s domain (A) eukaryotes, (B) bacteria, or (C) archaea.

Among the four archaea with missing CAT-anticodon tRNAs, only one appears to be due to a true isotype model misclassification. *Caldivirga maquilingensis* contains an extra tRNA^Met^, one of which we believe is a missing tRNA^Ile2^ based on the Ile (instead of Ile2) covariance model giving the highest score. For two other archaea, the absence of tRNA^Ile2(CAU)^ in *Nanoarchaeum equitans* and *Korarchaeum cryptofilum* is due to the presence of an uncommon tRNA^Ile3(UAU)^ obviating the need for the Ile2 tRNA, as explained in detail in previous studies (80). The remaining archaeon, *Methanosarcina mazei* Tuc01, with a missing tRNA^Ile2^ also does not have five other expected tRNAs (tRNA^Gly(UCC)^, tRNA^Gln(UUG)^, tRNA^Ser(UGA)^, tRNA^Thr(GGU)^, and tRNA^Leu(GAG)^), suggesting an incomplete genome assembly. Thus, the rate of misclassified CAT tRNAs is estimated to be 0.4% (1/217) for the analyzed archaeal genomes.

We found that over a hundred bacterial genomes lacked at least one tRNA^fMet^, tRNA^Met^, or tRNA^Ile2^ gene prediction, but for different reasons (Supplementary File S5). Similar to *Nanoarchaeum equitans* and *Korarchaeum cryptofilum* in Archaea, we discerned that seventeen bacterial genomes including species of the *Bifidobacterium, Mycoplasma*, and *Neorickettsia* genera have tRNA^Ile3(UAU)^ instead of tRNA^Ile2(CAU)^, with the majority previously identified (80,81). In addition, we found that 36 genomes with a missing fMet or Ile2 tRNA gene are caused by insertion of self-splicing group I introns (82,83) which tRNAscan-SE was not designed to detect. For example, previous studies determined the presence of a group I intron inserted at the anticodon loop of tRNA^fMet^ in *Gloeobacter violaceus* PCC 7421 and *Synechocystis sp*. PCC 6803 (82,83). We also manually identified group I introns in tRNA^Ile2^ of multiple *Acidovorax* and *Burkholderia* species by sequence alignments and RNA family similarity search (14,20). Aside from 43 genomes that may possibly have incomplete or erroneous assemblies and one with tRNA^Ile2^ split between the circular genome boundaries, the remaining 46 genomes have tRNA^Met^ misannotated as tRNA^Ile2^ or vice versa. These are apparently due to under-represented tRNA sequences in the consensus alignment used to generate the bacterial covariance models, particularly for divergent genera such as *Thermotoga* and *Desulfitobacterium*. Future versions of tRNAscan-SE may be able to address these outlier tRNAs by developing subdomain clade-specific models. However, given that only 1.2% (46/4036) of bacterial genomes have misclassified tRNA^Met^ or tRNA^Ile2^, this represents a massive improvement over the prior version of tRNAscan-SE.

### High confidence predictions for metazoan and plant genomes

Short interspersed repeated elements (SINEs) that are derived from tRNAs generally retain the well-conserved internal RNA polymerase III promoters found in tRNA genes, but often have mutations causing loss of typical cloverleaf secondary structure (33–35). This attenuated conservation pattern relative to true tRNAs enables covariance model analysis to identify them, in that their scores often fall below most true tRNA genes but are still detected above the default score threshold (20 bits) in the original version of tRNAscan-SE (1). tRNA-derived SINEs are numerous in mammalian genomes and some other large non-primate eukaryotes (33,84–87), resulting in many potentially inaccurate predictions for “young” SINE families that have retained most of the ancestral tRNA features. For example, the cat genome has over 403,500 tRNA predictions (the most among studied genomes) while the rat genome, previously reported to have many tRNA pseudogenes due to repetitive elements (84), has over 211,000 tRNA predictions (Table 4). Although tRNAscan-SE classifies over 80% of the predictions in these mammals as pseudogenes, the remaining still exceed our expected tally of roughly 400-600 tRNA predictions in vertebrate genomes that have few tRNA-derived SINE families. When comparing the non-pseudogene prediction score distributions between primates and other mammals, we noted that median scores in mammals such as cow and armadillo are significantly lower than those in primates like human (Supplementary Figure S6). In addition, plants like *Zea mays*, known to contain repetitive elements, also have lower prediction median scores. By checking against these repetitive elements well-studied in mouse (86), we found that a fair portion of the lowest-scoring tRNA predictions not categorized as pseudogenes are mostly part of the B1 or B2 repeats. Across different species, the inception of tRNA-derived repetitive elements is variable, leading to more or less sequence divergence from an ancestral tRNA, making hard cutoffs to remove repetitive elements non-trivial. For example, tRNA-derived SINES detected by tRNAscan-SE in marine mammals such as minke whale and dolphin tend to have relatively high tRNA scores (both the domain-specific eukaryotic model and isotype-specific), but can be identified because of a loss of normal base pairing in secondary structure (detectable by a drop in the tRNAscan-SE secondary structure score). We therefore developed a filtering tool that can be optionally applied to tRNAscan-SE results for better discrimination of functional tRNA genes. It assesses the predictions with a combination of domain-specific, isotype-specific, and secondary structure scores in two filtering stages after the pseudogene classification (see Methods), and determines the “high confidence” set of genes that are more strictly enriched for tRNAs likely capable of protein translation. By applying this filter, we are able to remove thousands of genes not marked as pseudogenes, but are clearly repetitive elements upon examination. For example, just over 4,000 additional repetitive elements in mouse (mean score 27.8 bits), and almost 17,000 in minke whale (mean score 48.0 bits) are identified and excluded from the high confidence set, allowing the core sets of just over 400 tRNA genes to emerge from the repetitive element noise. We recommend application of the high confidence filter for metazoans and plants which include the bulk of the tRNA-derived SINE-rich species we have tested. This quality filter may still allow a small number of tRNA-derived SINES into the high confidence set, as being overly aggressive risks the removal of unusual but potentially functional tRNA genes. While unicellular eukaryotes also sometimes contain tRNA-derived repetitive elements, the metazoan/plant filtering thresholds appear to remove numerous unusual but legitimate tRNAs. A different set of filter cutoffs (specified by the user in run options) would need to be developed and tested. tRNA-derived repetitive elements have not been found in bacteria or archaea, so this filter is not needed (and not recommended) for those species.

**Table 4.** tRNA predictions and post-filtered high confidence set in eukaryotic genomes with numerous repetitive elements. The top ten genomes with very large numbers of raw predictions are shown in comparison with human and mouse. High confidence predictions are determined as a result of the three-stage post-filtering process. The values in the table represent the number of gene predictions in each set, or removed by each filter.

| Genome | All tRNA predictions | tRNAscan-SE predicted pseudogenes | Predictions removed by secondary filter | Predictions removed by tertiary filter | High confidence set |
| --- | --- | --- | --- | --- | --- |
| Human | 596 | 85 | 91 | 0 | 417 |
| Mouse | 40,912 | 36,415 | 4,087 | 2 | 401 |
| Rat | 211,167 | 198,126 | 12,605 | 51 | 364 |
| Squirrel | 165,970 | 146,252 | 19,221 | 85 | 349 |
| Cat | 403,590 | 392,312 | 10,552 | 126 | 549 |
| Cow | 263,431 | 232,442 | 29,617 | 562 | 593 |
| Sheep | 256,819 | 225,920 | 29,710 | 420 | 534 |
| Ferret | 182,506 | 179,919 | 2,167 | 40 | 360 |
| Panda | 187,373 | 183,250 | 3,548 | 143 | 398 |
| Ferret | 182,506 | 179,919 | 2,167 | 40 | 360 |
| Minke whale | 212,492 | 194,977 | 13,916 | 2,879 | 428 |
| Dolphin | 172,909 | 157,354 | 12,694 | 2,236 | 368 |
| Armadillo | 227,726 | 154,522 | 72,576 | 89 | 462 |

An additional component of the high-confidence filter flags predictions that have high scores but atypical features such as unexpected anticodons (e.g., Pro-GGG, Asp-ATC, Tyr-ATA and 11 others in eukaryotes), or if they have a conflict of tRNA isotype identity based on comparison of the anticodon to the body of the tRNA (detailed in the next section). Because these types of atypical tRNAs are unusual and probably not functional but cannot be automatically assumed to be pseudogenes, they are specially marked for further investigation. In our study, the high confidence set remains below 1,000 tRNA genes in most genomes (Table 4), which provides researchers with a much better-enriched tRNA set to study and focus their experimental efforts.

### Isotype-specific covariance models improve functional annotation of tRNAs

Besides properly distinguishing the different CAU subtypes, isotype-specific covariance models have the potential to better classify or flag other unusual tRNA predictions. Among the genomes we studied, over 95% of the typical predicted tRNA genes have anticodons that match with the highest scoring isotype-specific model (Supplementary File S6). When inspecting cases in which the highest scoring isotype model and anticodon disagree, we surmised that the conflict could be caused for various reasons, including the possibility of altering the standard genetic code which has precedent (88). In our analyses of human tRNAs, the high-scoring gene tRNA-Leu-CAA-5-1 (66.5 bits) scores much higher with the tRNA^Met^ model than tRNA^Leu^ model (98.4 bits vs 2.6 bits). Its secondary structure (Figure 4B) shows the lack of a long variable arm, a typical identity element for tRNA^Leu(CAA)^ (Figure 4C), and it only has six nucleotide differences from tRNA-Met-CAT-3-1 (Figure 4A, Supplementary Figure S9A). The gene locus is present in most primates, yet only human, chimp, and gorilla have an unexpected A36 while the other genomes have U36 that would make it a typical tRNA^Met(CAU)^ (Supplementary Figure S9B). This suggests that the mutation was acquired relatively recently. Small RNA sequencing data from ARM-Seq (74), DM-tRNA-seq (75,76), and DASHR (77) studies show very low abundance level (4.98, 1.55, and 28 RPM respectively), indicating that this gene may no longer be expressed constitutively in human cells and is either a silenced pseudogene, or may conceivably have an alternate role under special regulation. The new high confidence filter identifies all cases of anticodon / isotype model conflicts, and annotates them when it is applied.

**Figure 4.**
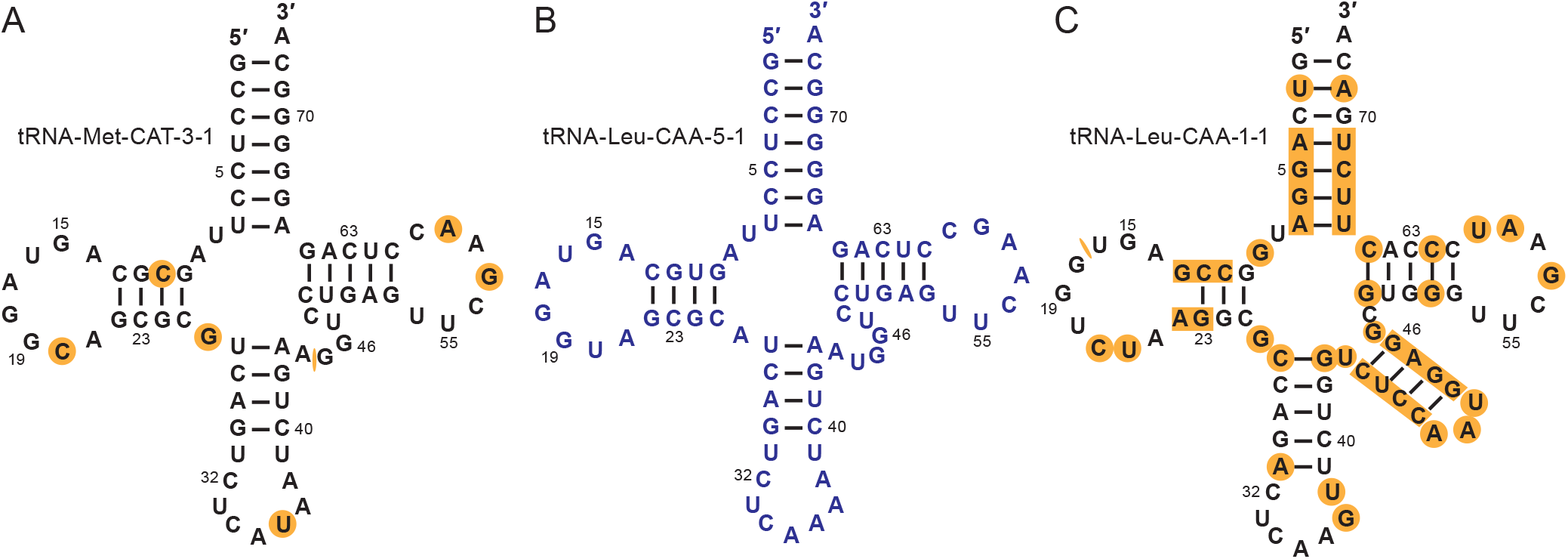
Isotype uncertainty in human tRNA-Leu-CAA-5-1. The primary sequence and the secondary structure comparison of (A) tRNA-Met-CAT-3-1, (B) tRNA-Leu-CAA-5-1, and (C) tRNA-Leu-CAA-1-1 shows that tRNA-Leu-CAA-5-1 is more similar to a tRNA^Met^ than a tRNA^Leu^ even though it has an anticodon that would decode leucine. The bases highlighted in orange represent the differences between the respective sequence and tRNA-Leu-CAA-5-1. The numbers next to the bases represent the Sprinzl canonical tRNA positions (16).

When reviewing fungal genomes, we found an unusual statistic for the multicellular fungus *Serpula lacrymans* (70): 70% of its non-pseudogene tRNA predictions showed isotype model / anticodon disagreement. Closer inspection revealed that the majority of these uncertain tRNA genes displayed three characteristics that are indicative of pseudogenes: (1) they are predicted to be tRNA^Ser^ based on the UGA anticodon, but the highest scoring isotype model is tRNA^Thr^; (2) the secondary structure of these predictions shows a variable loop instead of the expected variable arm which is an identity element for tRNA^Ser^; (3) the sequences of these high copy number elements are identical, and most of them have identical 50 bp upstream and downstream flanking regions, something not observed for authentic tRNA genes in fungi. This strongly suggests that they are repetitive elements and illustrates an example of using the anticodon / isotype model comparison to distinguish real tRNAs from previously unidentified repetitive elements. While this species is not recommended for application of the provided high confidence filter thresholds, the anticodon / isotype model conflict is annotated as “IPD” (Isotype Model Disagreement) whenever the “--detail” option is selected for tRNAscan-SE 2.0 runs.

Alternatively, isotype-and domain-specific classification may uncover lateral tRNA gene transfer events. We found a novel example for the archaeal methanogen *Methanobrevibacter ruminantium*, which has a tRNA^Arg^ with anticodon ACG that is not found in its closest sequenced relatives in the same genus, *Methanobrevibacter smithii* and *Methanobrevibacter sp*. AbM4. This is unusual, as A34 in tRNA^Arg^ is generally only found in bacteria and eukaryotes but not archaea, which use G34-containing tRNAs to decode pyrimidine-ending codons (67). Furthermore, the highest scoring isotype model prediction is for tRNA^Trp,^ which also disagrees with the Arg-ACG anticodon encoded in the gene. In a previous genome analysis, over 13% (294 out of 2217) of the coding genes in the bovine gut microbe *M. ruminantium* were identified to have originated from other species present in the same environment, including those from bacteria and eukaryotes (89). Horizontal gene transfer commonly occurs in microbes that share the same or similar habitats (90,91), yet most research has been focused on protein-coding genes. The predicted tRNA^Arg(ACG)^ scores 53.4 bits for the bacterial model and 55.1 bits for the eukaryotic model, in contrast to a score of 37.6 bits using the archaeal model. In addition, the highest scoring bacterial and eukaryotic isotype-specific models are both consistent with the anticodon, further supporting that this gene has been transferred from a species in another domain of life.

### Improved identification of archaeal tRNA genes with noncanonical introns

Some tRNAs in eukaryotes and archaea have introns that are removed by tRNA splicing endonuclease during maturation. The majority of archaeal tRNA introns are located one nucleotide downstream of the anticodon (position 37/38), yet some have been found at seemingly random, “noncanonical” positions in tRNA genes (5,36,68,92). In contrast to eukaryotic tRNA genes which are limited to one intron, archaeal tRNA genes can contain up to three, with *Pyrobaculum calidifontis* having the most tRNA introns (71 introns in 46 tRNA genes) out of all assessed complete genomes (36). These noncanonical introns have presented a challenge for predicting archaeal tRNA genes correctly. The introns preserve a general bulge-helix-bulge (BHB) secondary structure (68) that can be modelled using Infernal (20) for a covariance model search. In the previous version of tRNAscan-SE, we included a search routine and a covariance model to detect the noncanonical introns in archaea. The model was built using the known intron sequences at the time, mostly identified in Crenarchaeota. Due to slower performance of the previous Infernal versions, we included this routine as an optional feature to avoid significant increase to default search time. In addition, the original covariance model cannot effectively detect noncanonical tRNA introns in genomes that are more distantly related from those used for the training data. Therefore, we have redesigned the search process (see Methods; Supplementary Figure S3) by including two covariance models: one further optimized from the existing Crenarchaeal-focused model, and the other newly trained with introns from the archaeal phylum Thaumarchaeota. Both models take advantage of Infernal’s newest performance improvements, making it feasible to use both for the default archaeal search mode.

Among the 217 archaeal genomes analyzed, we found 1,533 canonical and 673 noncanonical introns in a total of 10,372 predicted tRNA genes (Supplementary Table S6). Previous study results and manual inspection show that our process has a low error rate, with just 2.9% of the noncanonical introns not detected due to extremely low similarity to the consensus, plus four noncanonical introns mis-annotated as canonical introns. Almost 70% of the identified introns across all studied species are canonical, while a small number of clades have more noncanonical than canonical tRNA introns. As described in previous studies, many tRNA genes in Thermoproteales species (a family within the Crenarchaeota) are known to harbor multiple introns (36,37). In our analysis, we found an average of 0.84 intron per tRNA gene in Thermoproteales while the overall average in the studied archaeal genomes is 0.21 per gene. In comparison with tRNAscan-SE 1.3, the new search process identifies 118 (21%) additional noncanonical introns with 54 of them previously mis-annotated as canonical introns. Thaumarchaeota genomes that were originally predicted to have a total of 58 noncanonical introns are now found to harbor 107 noncanonical introns. This shows a significant improvement to accurately identify archaeal tRNA introns with the use of the clade-specific covariance models.

### Predicting mitochondrial tRNAs in vertebrates

Mitochondrial genomes in vertebrates, representing the majority of available sequenced mitochondrial genomes, typically include 22 tRNA (mt-tRNA) genes (93). Each mt-tRNA decodes one of the twenty amino acids, except for leucine and serine which are decoded by two mt-RNAs with distinct anticodons. Previous studies have shown that sequence and structural simplification has resulted in mt-tRNAs to commonly deviate from the canonical clover-leaf structure of their ancestral bacterial tRNAs (94). In addition to abbreviated stem and loop lengths in D-and T-arms, complete loss of the D-arm is also observed in most vertebrate mt-tRNA^Ser(GCU)^ (66).

The original tRNAscan-SE was not designed to identify mitochondrial tRNAs, but it offered an ad hoc “organellar search mode” which employed a general covariance model trained with a mix of eukaryotic cytosolic, bacterial, and archaeal tRNAs, plus a more permissive detection cutoff. As such, its performance was mediocre, and in particular it misses mt-tRNAs with highly degenerate secondary structure. For example, in mammals, it consistently fails to detect the mt-tRNA^Ser(GCU)^ which lacks the D-arm (29). To effectively identify mt-tRNAs in vertebrates, we developed 22 new covariance models, one for each specific isotype/anticodon. tRNAscan-SE 2.0 scans input sequences with all 22 mt-tRNA covariance models, merges results, and annotates each detected tRNA locus with the identity and score of the highest scoring model.

To assess the false positive rate of mt-tRNA gene prediction, we applied the new vertebrate mitochondrial search mode to a negative test set of 17,500 virtual mitochondrial genomes generated with varying G/C content (Supplementary Table S4 and Supplementary File S4). For tRNAscan-SE 2.0, no false positives were found at or above the default score cutoff (20 bits) within the virtual genomes, which confirms very high specificity characteristic of prior tRNA covariance models at that score threshold (Supplementary Table S4, Supplementary Figure S2B and S4, Supplementary File S8). We then applied existing mitochondrial search tools MiTFi (9) and ARWEN (8) to the same set of virtual genomes for comparison of false positive rates (Supplementary Table S4). Similar to tRNAscan-SE 2.0, MiTFi did not have any false positives below its recommended e-value. In contrast, ARWEN predicted 34,634 false positive hits with 11,949 (34.5%) classified as possible pseudogenes, or about 2 to 3 false positives per virtual genome, which is consistent with a relatively high false positive rate as previously reported (8).

We assessed sensitivity by comparing prediction results from a positive test set of 3,345 actual mitochondrial genomes from vertebrates (Supplementary File S7) which contain 73,674 annotated mt-tRNA genes in NCBI RefSeq (24) (Table 5, Supplementary File S9). Predictions consistent with RefSeq annotations were similarly high for tRNAscan-SE 2.0 (99.0%) and MiTFi (98.8%). ARWEN was significantly lower at only 87.7% consistency with RefSeq annotations, meaning more than one in ten predictions have no support in existing annotations. Regarding execution speed, tRNAscan-SE 2.0 and ARWEN performed similarly at 5.5 and 4.5 seconds per mitochondrial genome (average ∼3 Kbp/sec) (Table 5). The processing time of MiTFi was almost 60 times longer (average ∼50 bp/sec), although this is likely due to the use of an older version of the Infernal search engine – if it were updated to the newest version, we expect it would perform similarly to tRNAscan-SE 2.0.

**Table 5.** Comparison of mt-tRNA gene predictions with NCBI RefSeq annotations. 73,674 annotated mt-tRNAs in 3,345 mitochondrial genomes from NCBI RefSeq (24) were used as references for comparison. “Consistent” represents the count and percentage of predicted mt-tRNA genes that are the same or nearly the same as RefSeq annotations (may include up to a 15-nt difference in start and/or end positions). “Isotype mismatch” represents the number and percentage of predicted mt-tRNA genes that have conflicting isotype classification. “Position mismatch” represents the number and percentage of predicted mt-tRNA genes that had conflicting strand or start/end positions beyond +/-15 nts. “Novel” represents the predicted mt-tRNA genes that were not found in RefSeq annotations. “Not detected” is the count of RefSeq annotations that were not detected by the prediction algorithm.

| Prediction algorithm | Predicted mt-tRNAs | Consistent | Isotype mismatch | Position mismatch | Novel | Not detected |
| --- | --- | --- | --- | --- | --- | --- |
| tRNAscan-SE 2.0 (vertebrate mito mode) | 73,671 | 72,920 (99.0%) | 35 (0.05%) | 527 (0.7%) | 189 (0.3%) | 192 |
| ARWEN | 80,617 | 70,677 (87.7%) | 211 (0.3%) | 551 (0.7%) | 9,178 (11.4%) | 2,235 |
| MitFi | 73,685 | 72,796<br>(98.8%) | 96 (0.1%) | 528 (0.7%) | 265 (0.4%) | 254 |

Although NCBI RefSeq (24) has a well-established collection of mitochondrial genomes, most of the mt-tRNA genes were annotated with a combination of “imperfect” gene prediction tools such as tRNAscan-SE 1.3 organellar mode, BLAST (95) sequence similarity search, and manual curation, leading to possible misannotations. For example, when inspecting the tRNAscan-SE 2.0 predictions that have position inconsistencies with RefSeq annotations, we found that over 99% of them (522 out of 527) have identical sequence boundaries but with opposite strandedness. Since the predictions from the other two tools also present this comparison discrepancy, it suggests that those genes in RefSeq were annotated with the incorrect strand. Similarly, over 70% of the predictions that have conflicting isotypes (25 out of 35) were found to have incorrect isotype classifications in RefSeq by switching between mt-tRNA^Asn^ and mt-tRNA^Asp^ or between mt-tRNA^Gln^ and mt-tRNA^Glu^. The remaining disagreements are caused by unexpected anticodons that require further studies. In addition, conflicting genomic coordinates in RefSeq gene annotations, such as identical sequence boundaries of mt-tRNA^Ala^ and mt-tRNA^Trp^ in *Solenostomus cyanopterus*, results in both novel and missing predictions in the comparison. On the other hand, gene order rearrangements caused by models such as tandem duplication and random loss (TDRL) occur in some vertebrate mitochondrial genomes and may lead to duplication of mt-tRNA genes (9,96–98). For instance, a cluster of extra-copy mt-tRNA genes decoding tyrosine, cysteine, asparagine, and alanine were detected in *Diretmus argenteus* by tRNAscan-SE 2.0 and MiTFi, suggesting a pattern of tandem duplication that has not been previously reported. With the absence of many of the duplicated mt-tRNA genes in RefSeq annotations, these predictions from tRNAscan-SE 2.0 contribute to the majority of the novel predictions in the RefSeq comparison.

Given the combination of lowest false positivity rate, highest sensitivity rate, and fast execution, we suggest tRNAscan-SE 2.0 as the new standard for vertebrate mitochondrial tRNA genome analyses, and it could be applied to correct years of accumulated errors or omissions in public database annotation.

## DISCUSSION

tRNAscan-SE 2.0 has been in development, testing and refinement for over five years, and is the culmination of an improved search engine (Infernal v1.1), streamlined search strategy, updated and expanded search models, improved isotype predictions (fMet/iMet/Met/Ile2, SeC), new features (archaeal multi-intron prediction, high confidence set prediction), annotation of atypical tRNAs (unexpected anticodons, anticodon/isotype model conflicts) and new targets (vertebrate mitochondrial genomes). The 2.0 program functionality has been available for web user analyses during this time (http://trna.ucsc.edu/tRNAscan-SE/) (3), and currently averages over 1,100 users per month. It has also been vetted and put into service at major genome centers such as the latest version of NCBI Prokaryotic Genome Annotation Pipeline (99), DOE-JGI IMG annotation pipeline (https://img.jgi.doe.gov/docs/pipelineV5/), and RNA 2D Templates (R2DT) at RNAcentral for RNA secondary structure visualization (100), with the 2.0 source code package downloaded over 18,000 times (Bioconda, GitHub, and Lowe Lab website). The results from tRNAscan-SE 2.0 have been publicly available in GtRNAdb 2.0 (5) and RNAcentral (101,102) for over three years. Here, we have thoroughly documented all new features and critically assessed the program’s performance on more than eight thousand complete or nearly complete genomes.

The multi-isotype model strategy implemented by tRNAscan-SE 2.0 provides an essential new perspective for identification of functional ambiguity of likely pseudogenes as well as atypical tRNA genes worthy of further investigation. “Hybrid” tRNAs with anticodons that do not match the expected isotype sequence features can now be detected, providing examples of tRNAs which could alter the genetic code in special situations such as cellular damage or other stress conditions (103,104). While we have not biochemically verified the expression or efficiency of these candidate “genetic code-breakers”, our tool can readily reveal them and invite investigation by the community. In other cases, anticodon-isotype model disagreement are strong indicators of tRNA-derived repetitive elements which are slowly decaying, thus anticodon mutations affecting codon translation fidelity are not selected against. We have used this frequent property of non-functional tRNA-derived elements plus new secondary structure score thresholding to greatly reduce repetitive element noise, producing “high confidence” sets that are better enriched for functional core tRNA genes in metazoans and plants. This should help researchers prioritize the most biologically relevant tRNA genes for genetic manipulation using CRISPR and other powerful tools to dissect individual tRNA gene regulation and function. For example, the mouse and rat tRNA sets now contain 401 and 364 high confidence tRNA genes, respectively, versus the prior tRNAscan-SE 1.3 analysis tallies of ∼4,400 tRNAs and ∼13,000 respective non-pseudogenes.

New models for proper classification and annotation of CAU-anticodon tRNAs into initiator methionine, elongator methionine, and isoleucine-2 groups are long overdue, and will improve the functional annotation for 5-10% of all tRNAs. With detection of selenocysteine tRNAs greatly improved, research in the regulation of selenoprotein production will be better supported. Mitochondrial tRNAs have also been overlooked for over twenty years, and now with tRNAscan-SE 2.0’s new set of mitochondrial models, it may become the new reference tool for uniform, reliable vertebrate mitochondrial genome analyses.

With this update, researchers can now detect archaeal tRNA genes with noncanonical tRNA introns, by default, in addition to those with canonical introns. Although this feature was first introduced in v1.3, the initial intron-detecting covariance model and unoptimized processing resulted in a significant increase of execution time for archaeal genomes, and in some cases, a high rate of missing introns affecting as many as one-third of tRNA genes per genome. With newly created intron models enabling detection of two distinct intron types, and complete redesign of the intron search process, the new release provides comprehensive identification of both canonical and noncanonical tRNA introns. Also, due to misidentification of archaeal introns by v.1.3, it sometimes caused incorrect anticodon and tRNA isotype prediction; v2.0 consequently improves accuracy of archaeal tRNA functional annotation as well.

In spite of the multitude of improvements to tRNA detection, additional enhancements and capabilities are needed. Vertebrate mitochondrial genomes were a practical first target for organellar analyses due to their biomedical relevance and their consistent, mostly canonical tRNA secondary structures. We plan to broaden this capability to mt-tRNA detection in insects, nematodes, fungi, and other heavily studied clades. Some mt-tRNAs in these clades have evolved to contain highly degenerate forms missing one or two tRNA arms (105–107), and therefore, more diverse model sets will be needed to attain high sensitivity and annotation accuracy. We anticipate new model sets can be developed by or in collaboration with specific user communities, and we welcome these joint projects. We also believe more work can be done to further improve functional annotations and better classify true tRNA-derived SINEs / pseudogenes / silenced tRNA genes versus those actively transcribed and/or likely used in translation. For these goals, we will look to integrating contextual evolutionary information predicting gene activity (108), contextual genomic information (e.g., flanking regions characteristic of repetitive elements, transcription termination signals), or other types of readily available high throughput sequencing data (e.g., tRNA-seq, ribosome profiling, ATAC-seq).

Furthermore, more specialized models are needed for non-organellar tRNAs as we find more exceptional tRNAs in the tree of life. For example, the new isotype-specific models occasionally do not give the highest score for the true isotype identity, most likely due to exceptional features in specific isotypes of some species. Thus, we believe more clade-specific isotype-specific models will be needed to accurately represent the true variation found in nature. Why this evolutionary variation is so widespread is not easily explained by tRNA’s well-defined, relatively unchanging role in protein translation. Fairly recent discoveries of alternate regulatory roles of tRNAs, and in particular tRNA-derived small RNAs (tDRs), may reveal the impetus for unexpected tRNA sequence variation, requiring new models to identify tRNA subtypes with specialized functions. Characterizing the precise functionality, if any, associated with the growing gallery of atypical / borderline tRNAs that show experimental or evolutionary evidence of biological importance is beyond the scope of this work, but we anticipate there are myriad new discoveries primed by tRNAscan-SE 2.0 improved annotation.

Together with the ever-growing pool of genomes, the continued enhancement of tRNAscan-SE should assure its role as an essential tool for expanding the world of tRNA biology.

## DATA AVAILABILITY

tRNAscan-SE is open source and released under GNU General Public License v3. The installation package with source code can be downloaded from GitHub repository https://github.com/UCSC-LoweLab/tRNAscan-SE and http://trna.ucsc.edu/tRNAscan-SE/.

## SUPPLEMENTARY DATA

Supplementary data are available at NAR online.

## ACKNOWLEDGEMENT

We would like to thank Aaron Cozen for his valuable feedback during the development of tRNAscan-SE 2.0. We also thank our lab members for their comments on software functionality and the manuscript.

