## Supplemental Tables and Figures for "tRNAscan-SE 2.0: Improved Detection and Functional Classification of Transfer RNA Genes"

### SUPPLEMENTARY TABLES AND FIGURES

**Supplementary Table S1** Data source of specific genomes analyzed in this manuscript.

| Genome | Domain | Assembly/Accession |
|---|---|---|
| *Arabidopsis thaliana* | Eukaryota | TAIR10 |
| Armadillo (*Dasypus novemcinctus*) | Eukaryota | Dasnov3.0 |
| Budding yeast (*Saccharomyces cerevisiae* S288C) | Eukaryota | GCA_000146045.2/R64 |
| *Caenorhabditis elegans* | Eukaryota | WBcel235 |
| Cat (*Felis catus*) | Eukaryota | Felis_catus_9.0 |
| Chicken (*Gallus gallus*) | Eukaryota | GRCg6a |
| Cow (*Bos taurus*) | Eukaryota | Bos_taurus_UMD_3.1.1 |
| Dolphin (*Tursiops truncates*) | Eukaryota | Ttru_1.4 |
| Elephant shark (*Callorhinchus milii*) | Eukaryota | Callorhinchus_milii-6.1.3 |
| Ferret (*Mustela putorius furo*) | Eukaryota | MusPutFur1.0 |
| Human (*Homo sapiens*) | Eukaryota | GRCh37/hg19 and GRCh38/hg38 |
| Japanese rice (*Oryza sativa* Japonica Group) | Eukaryota | IRGSP-1.0 |
| Maize (*Zea mays*) | Eukaryota | RefGen_v4 |
| Medaka (*Oryzias latipes*) | Eukaryota | GCA_002234675.1/ASM223467v1 |
| Minke whale (*Balaenoptera acutorostrata scammoni*) | Eukaryota | BalAcu1.0 |
| Mouse (*Mus musculus*) | Eukaryota | GRCm38/mm10 |
| Panda (*Ailuropoda melanoleuca*) | Eukaryota | AilMel_1.0 |
| Rat (*Rattus norvegicus*) | Eukaryota | Rnor6.0 |
| Rhesus (*Macaca mulatta*) | Eukaryota | Mmul8.0.1 |
| Sheep (*Ovis aries*) | Eukaryota | Oar_v4.0 |
| Squirrel (*Spermophilus tridecemlineatus*) | Eukaryota | SpeTri2.0 |
| *Bacillus subtilis* subsp. subtilis str. 168 | Bacteria | GCA_000009045.1/ASM904v1 |
| *Clostridioides difficile* 630 | Bacteria | GCA_000009205.2/ASM920v2 |
| *Escherichia coli* str. K-12 substr. MG1655 | Bacteria | GCA_000005845.2/ASM584v2 |
| *Gloeobacter violaceus* PCC 7421 | Bacteria | GCA_000011385.1/ ASM1138v1 |
| *Mycobacterium tuberculosis* H37Rv | Bacteria | GCA_000195955.2/ASM19595v2 |
| *Pseudomonas aeruginosa* PAO1 | Bacteria | GCA_000006765.1/ASM676v1 |
| *Synechocystis sp.* PCC 6803 | Bacteria | GCA_000009725.1/ ASM972v1 |
| *Halobacterium sp.* NRC-1 | Archaea | GCA_000006805.1/ASM680v1 |
| *Methanobrevibacter ruminantium* M1 | Archaea | GCA_000024185.1/ASM2418v1 |
| *Methanobrevibacter smittii* ATCC 35061 | Archaea | GCA_000016525.1/ASM1652v1 |
| *Methanobrevibacter sp.* AbM4 | Archaea | GCA_000404165.1/ASM40416v1 |
| *Methanocaldococcus jannaschii* DSM 2661 | Archaea | GCA_000091665.1/ASM9166v1 |
| *Pyrobaculum aerophilum* IM2 | Archaea | GCA_000007225.1/ASM722v1 |
| *Pyrobaculum calidifontis* JCM 11548 | Archaea | GCA_000015805.1/ASM1580v1 |
| *Pyrococcus furiosus* DSM 3638 | Archaea | GCA_000007305.1/ASM730v1 |
| *Sulfolobus acidocaldarius* DSM 639 | Archaea | GCA_000012285.1/ASM1228v1 |
| *Chirocentrus dorab* mitochondrion | Eukaryota | NC_006913.1 |
| *Desmognathus fuscus* mitochondrion | Eukaryota | NC_006339.1 |
| *Diretmus argenteus* mitochondrion | Eukaryota | NC_008127.1 |
| *Hemiechinus auritus* mitochondrion | Eukaryota | NC_005033.1 |
| *Homo sapiens* mitochondrion | Eukaryota | NC_012920.1 |
| *Solenostomus cyanopterus* mitochondrion | Eukaryota | NC_010267.1 |
| *Taeniopygia guttata* mitochondrion | Eukaryota | NC_007897.1 |

**Supplementary Table S2** tRNAscan-SE performance comparison for non-default "maximum sensitivity" mode. Nine genomes representing different clades were benchmarked on COVE-only and maximum sensitivity search mode using tRNAscan-SE 1.3 and new version 2.0 respectively. The genome scans using tRNAscan-SE 2.0 were conducted with Infernal (1) running eight threads in parallel. (A) Total number of tRNA predictions and the number of predictions excluding those determined as possible pseudogenes in the nuclear genomes are listed. (B) Reported search times are based on using a server with dual Intel 10-core HT processors at 2.30GHz. tRNAscan-SE 1.3 only allows processing with a single thread while tRNAscan-SE 2.0 makes use of the multi-core processor architecture for parallel computation. *The processing time using tRNAscan-SE 1.3 COVE-only mode for human, mouse, and maize was estimated with the result processing time of chromosomes 17, 13, and 10 in respective genomes. The total number of tRNA predictions for these genomes are therefore not available (NA).

**A**

| Genome | Domain | tRNAscan-SE 1.3 | | tRNAscan-SE 2.0 | |
|---|---|---|---|---|---|
| | | Total tRNA predictions | tRNAs excluding pseudogenes | Total tRNA predictions | tRNAs excluding pseudogenes |
| *Escherichia coli* K-12 | Bacteria | 88 | 87 | 90 | 88 |
| *Pyrococcus furiosis* DSM 3638 | Archaea | 46 | 46 | 46 | 46 |
| *Saccharomyces cerevisiae* | Eukaryota | 275 | 275 | 275 | 275 |
| *Homo sapiens* | Eukaryota | NA | NA | 626 | 516 |
| *Mus musculus* | Eukaryota | NA | NA | 41,942 | 4,165 |
| *Caenorhabditis elegans* | Eukaryota | 837 | 620 | 724 | 618 |
| *Drosophila melanogaster* | Eukaryota | 297 | 292 | 295 | 292 |
| *Arabidopsis thaliana* | Eukaryota | 648 | 635 | 643 | 633 |
| *Zea mays* | Eukaryota | NA | NA | 2,103 | 1,392 |

**B**

| Genome | Domain | tRNAscan-SE 1.3 | tRNAscan-SE 2.0 |
|---|---|---|---|
| *Escherichia coli* K-12 | Bacteria | 9h 33m | 15m 18s |
| *Pyrococcus furiosis* DSM 3638 | Archaea | 3h 52m | 14m 43s |
| *Saccharomyces cerevisiae* | Eukaryota | 1d 1h | 1h 1m |
| *Homo sapiens* | Eukaryota | 94d 12h* | 5d |
| *Mus musculus* | Eukaryota | 175d 19h* | 5d 9h |
| *Caenorhabditis elegans* | Eukaryota | 7d 3h | 5h 13m |
| *Drosophila melanogaster* | Eukaryota | 9d 11h | 9h 18m |
| *Arabidopsis thaliana* | Eukaryota | 8d 12h | 6h 4m |
| *Zea mays* | Eukaryota | > 110d* | 4d 12h |

**Supplementary Table S3** tRNA predictions for virtual genomes. Virtual genomes were generated using a $5^{th}$ order Markov chain trained from the original genome. tRNAscan-SE 2.0 default mode with a score cutoff of 10 bits (10 bits below default cutoff) was used to scan the virtual genomes. The number of tRNA hits represents the predictions identified in all virtual genomes of the original one that score 10 bits or above.

| Genome | Domain | GC content | # of Virtual Genomes | # of tRNA Hits | Score Range (bits) |
|---|---|---|---|---|---|
| *Escherichia coli* K-12 | Bacteria | 50.8% | 100 | 37 | 10.0 – 16.2 |
| *Halobacterium sp.* NRC-1 | Archaea | 67.9% | 100 | 39 | 10.0 – 19.8 |
| *Saccharomyces cerevisiae* S288C | Eukaryota | 38.4% | 100 | 1 | 16.9 |
| *Homo sapiens* | Eukaryota | 42.4% | 100 | 610<br>17 | 10.0 – 19.8<br>20.2 – 29.5 |

**Supplementary Table S4** Mitochondrial tRNA predictions for virtual genomes. tRNAscan-SE 2.0 mitochondrial mode with score cutoff set to 0 bits (20 bits below default cutoff) was used to scan the virtual genomes. The number of tRNAscan-SE hits represents the predictions identified in all virtual genomes that scored 0 bits or above. The number of tRNA hits from ARWEN include total predictions across all virtual genomes with predicted non-pseudogenes in parentheses. MiTFi (3) default search mode was used, showing predictions identified in all virtual genomes, with none more significant than the suggested e-value cutoff (0.0001) in parentheses.

| Mitochondrial Genome | GC content | # of Virtual Genomes | tRNAscan-SE 2.0 | | ARWEN | MiTFi |
|---|---|---|---|---|---|---|
| | | | # of tRNA Hits | Score Range (bits) | # of tRNA Hits (non-pseudogenes) | # of tRNA Hits (better than e-value cutoff) |
| *Homo sapiens* | 44.4% | 3,500 | 196 | 3.0 – 8.4 | 2,679 (1,245) | 3 (0) |
| *Hemiechinus auritus* | 30.9% | 3,500 | 275 | 0.3 – 16.5 | 3,908 (2,077) | 4 (0) |
| *Chirocentrus dorab* | 53.2% | 3,500 | 81 | 0.0 – 6.2 | 8,241 (6,344) | 0 (0) |
| *Desmognathus fuscus* | 31.0% | 3,500 | 624 | 1.2 – 9.8 | 12,096 (7,320) | 1 (0) |
| *Taeniopygia guttata* | 45.9% | 3,500 | 59 | 0.4 – 5.5 | 7,712 (5,701) | 0 (0) |

**Supplementary Table S5** Published human RNA-seq abundance for tRNAs predicted by one or both versions of tRNAscan-SE for human genome assembly GRCh37/hg19. (A) Predictions, excluding pseudogenes, only reported by tRNAscan-SE 1.3 and (B) predictions only reported by tRNAscan-SE 2.0 (default search mode) show very low or no detectable abundance based on prior small RNA sequencing studies employing ARM-Seq (4), DM-tRNA-seq (5), or collected by the database DASHR v2 (6), shown in reads per million (RPM). ARM-Seq and DM-tRNA-seq data were processed by using tRAX (https://github.com/UCSC-LoweLab/tRAX) as described in ARM-Seq publication (4). RPM was calculated as (raw read count aligned to a transcript / total number of alignments) x 1,000,000. The total number of alignments instead of unique reads was used in order to allow multiple mappings per read, as there are frequently multiple identical tRNA gene copies in the genome. The highest RPMs of the tissues/conditions in each study were reported. For comparison, (C) tRNA transcripts with data from all three studies predicted by both tRNAscan-SE 1.3 and 2.0, show clear, consistent evidence of transcription. RPM values above the minimum cutoff of 5.0 highlighted in bold.

**A**

| GtRNAdb ID | Locus | tRNAscan-SE 1.3 Score | ARM-Seq RPM | DM-tRNA-seq RPM | DASHR v2 RPM |
|---|---|---|---|---|---|
| tRNA-Asp-GTC-7-1 | chr1:161501915-161501986 (+) | 34.1 | 3.03 | 0.22 | **29.91** |
| tRNA-Gln-CTG-15-1 | chr9:126655522-126655594 (-) | 23.0 | 0.00 | 0.00 | 0.58 |
| tRNA-Gln-CTG-18-1 | chr16:71724890-71724963 (+) | 20.8 | 0.00 | 0.00 | 0.01 |
| tRNA-Glu-CTC-17-1 | chr1:149334272-149334340 (+) | 21.6 | 0.09 | 0.00 | 4.19 |
| tRNA-Glu-CTC-6-1 | chr18:43299751-43299822 (+) | 32.1 | **81.65** | **30.82** | 1.27 |
| tRNA-Glu-CTC-8-1 | chr3:125413177-125413248 (-) | 29.9 | 0.15 | 0.20 | 2.97 |
| tRNA-Glu-TTC-16-1 | chr1:149719802-149719870 (+) | 21.3 | 0.09 | 0.00 | 4.19 |
| tRNA-His-GTG-3-1 | chr3:148316563-148316634 (-) | 22.5 | 0.25 | 0.00 | 0.91 |
| tRNA-Leu-CAG-3-1 | chr5:159392041-159392118 (-) | 20.5 | 0.00 | 0.00 | 0.00 |
| tRNA-Lys-TTT-16-1 | chr19:19852207-19852277 (+) | 28.3 | 0.34 | 0.31 | 2.23 |
| tRNA-Sup-CTA-1-1 | chr17:15408685-15408758 (+) | 28.0 | 0.00 | 0.00 | 0.01 |
| tRNA-Sup-TTA-2-1 | chr21:15926516-15926586 (+) | 22.4 | 0.00 | 0.00 | 0.00 |
| tRNA-Trp-CCA-6-1 | chr9:115616989-115617087 (+) | 23.4 | 0.00 | 0.00 | 0.40 |
| tRNA-Trp-CCA-7-1 | chr11:45290200-45290273 (-) | 21.2 | 0.00 | 0.00 | 0.01 |
| tRNA-Val-CAC-12-1 | chr6:27650488-27650561 (+) | 33.4 | 0.14 | 0.10 | 1.05 |

**B**

| GtRNAdb ID | Locus | tRNAscan-SE 2.0 Score | ARM-Seq RPM | DM-tRNA-seq RPM | DASHR v2 RPM |
|---|---|---|---|---|---|
| tRX-Ala-NNN-1-1 | chr6:26713898-26713970 (+) | 24.8 | 0.00 | 2.33 | 0.12 |
| tRX-Ala-NNN-1-2 | chr6:26788156-26788228 (-) | 24.8 | 0.00 | 2.33 | 0.12 |
| tRX-Ala-NNN-2-1 | chr6:58156450-58156522 (-) | 26.3 | 0.00 | 2.33 | 0.12 |
| tRX-Ala-NNN-3-1 | chr19:12299376-12299449 (+) | 37.8 | 1.36 | 0.14 | 0.04 |
| tRX-Ala-NNN-4-1 | chr16:80479452-80479528 (-) | 38.2 | 0.00 | 0.00 | 0.23 |
| tRX-Asn-NNN-1-1 | chr1:147789485-147789554 (-) | 20.4 | 0.00 | 0.00 | **400.0** |
| tRX-Asp-NNN-1-1 | chr6:28795190-28795261 (-) | 34.9 | **19.03** | 3.62 | **12.27** |
| tRX-Cys-NNN-1-1 | chr7:149103155-149103226 (-) | 34.0 | 0.00 | 0.06 | 0.05 |
| tRX-Cys-NNN-2-1 | chr8:9939323-9939393 (-) | 27.2 | 2.52 | 0.07 | 4.38 |
| tRX-Cys-NNN-4-1 | chr7:149325244-149325326 (+) | 26.6 | 3.39 | 0.00 | **136.8** |
| tRX-Gln-NNN-1-1 | chr8:89145236-89145309 (-) | 21.5 | 0.00 | 0.00 | 0.00 |
| tRX-Glu-NNN-1-1 | chr8:11770800-11770871 (+) | 21.9 | **45.29** | 1.27 | **96.75** |
| tRX-Ile-NNN-1-1 | chrX:3833685-3833758 (-) | 46.5 | 0.00 | 0.00 | 0.03 |
| tRX-Ile-NNN-1-2 | chrX:3795256-3795329 (-) | 46.5 | 0.00 | 0.00 | 0.03 |

| GtRNAdb ID | Locus | tRNAscan-SE 2.0 Score | ARM-Seq RPM | DM-tRNA-seq RPM | DASHR v2 RPM |
|---|---|---|---|---|---|
| tRX-Ile-NNN-3-1 | chr6:27228710-27228784 (+) | 32.4 | 0.00 | 0.00 | 4.07 |
| tRX-Leu-NNN-2-1 | chr11:113432995-113433078 (-) | 45.5 | 0.00 | 0.28 | 0.24 |
| tRX-Leu-NNN-3-1 | chr2:152687652-152687719 (-) | 22.0 | 0.00 | 0.00 | 0.00 |
| tRX-Lys-NNN-1-1 | chr16:2977662-2977734 (+) | 46.8 | 0.85 | 0.50 | **33.70** |
| tRX-Lys-NNN-2-1 | chr3:15252973-15253048 (+) | 41.4 | 4.04 | 1.50 | **55.81** |
| tRX-Lys-NNN-6-1 | chr5:167816644-167816715 (-) | 29.9 | 0.00 | 0.17 | 4.48 |
| tRX-Ser-NNN-2-1 | chrX:25280089-25280160 (-) | 21.6 | 0.00 | 0.00 | 0.00 |
| tRX-Tyr-NNN-1-1 | chr14:21137333-21137418 (-) | 22.8 | **6.89** | 0.12 | **7.93** |
| tRX-Val-NNN-1-1 | chr6:158824648-158824720 (-) | 33.8 | **555.85** | **85.85** | **34.54** |
| tRX-Val-NNN-2-1 | chr2:85085876-85085948 (+) | 47.6 | **6.94** | 0.26 | **6.12** |

C

| GtRNAdb Transcript ID | ARM-Seq RPM | DM-tRNA-seq RPM | DASHR v2 RPM |
|---|---|---|---|
| tRNA-Arg-ACG-1 | **2516.19** | **3920.81** | **331.43** |
| tRNA-Arg-CCG-2 | **197.66** | **1687.58** | **8602.75** |
| tRNA-Arg-CCT-4 | **140.86** | **697.99** | **1570.98** |
| tRNA-Arg-TCG-3 | **353.59** | **2747.50** | **942.71** |
| tRNA-Arg-TCG-5 | **107.45** | **1353.45** | **76.28** |
| tRNA-Arg-TCT-1 | **166.02** | **761.73** | **3335.3** |
| tRNA-Asn-GTT-3 | **483.26** | **7733.13** | **353.23** |
| tRNA-Gln-CTG-1 | **8811.83** | **30231.34** | **546.83** |
| tRNA-Gln-TTG-1 | **839.62** | **1822.19** | **331.83** |
| tRNA-Gly-CCC-2 | **854.22** | **6031.06** | **19157.94** |
| tRNA-His-GTG-1 | **47604.36** | **47683.55** | **4234.54** |
| tRNA-iMet-CAT-1 | **4574.68** | **48294.91** | **953.67** |
| tRNA-Leu-AAG-1 | **388.43** | **1448.88** | **104.26** |
| tRNA-Leu-AAG-2 | **689.97** | **2625.32** | **244.66** |
| tRNA-Leu-CAA-1 | **436.53** | **7708.91** | **59.73** |
| tRNA-Leu-CAA-3 | **126.11** | **1583.25** | **69.99** |
| tRNA-Leu-CAA-4 | **316.77** | **1894.11** | **693.69** |
| tRNA-Leu-CAG-1 | **1073.59** | **16999.12** | **377.06** |
| tRNA-Leu-CAG-1 | **889.63** | **16999.12** | **377.06** |
| tRNA-Leu-CAG-2 | **368.80** | **4271.44** | **374.48** |
| tRNA-Leu-TAA-1 | **48.00** | **1570.88** | **1647.96** |
| tRNA-Leu-TAA-3 | 4.79 | **677.20** | **277.74** |
| tRNA-Leu-TAG-1 | **130.93** | **565.19** | **121.69** |
| tRNA-Leu-TAG-3 | **124.25** | **519.45** | **806.86** |
| tRNA-Lys-CTT-1 | **1169.83** | **3386.12** | **29182.64** |
| tRNA-Met-CAT-1 | **113.52** | **662.97** | **1276.43** |
| tRNA-Met-CAT-6 | **177.66** | **973.06** | **903.79** |
| tRNA-Phe-GAA-1 | **1116.33** | **23229.68** | **428.89** |
| tRNA-Pro-TGG-3 | **5872.17** | **14288.90** | **195.43** |
| tRNA-Ser-AGA-2 | **3985.58** | **17109.11** | **527.92** |
| tRNA-Ser-GCT-6 | **332.23** | **1366.31** | **1328.29** |
| tRNA-Ser-TGA-3 | **673.52** | **3012.75** | **526.39** |
| tRNA-Thr-TGT-4 | **338.12** | **1034.45** | **250.59** |
| tRNA-Val-AAC-1 | **3917.56** | **65305.70** | **5175.19** |

**Supplementary Table S6** Archaeal tRNA intron predictions by clade, type, and count per gene.

| Clade | # of Genomes | # of tRNA Genes | # of Canonical Introns | # of Non-canonical Introns | # of Introns Per tRNA Gene | | | |
|---|---|---|---|---|---|---|---|---|
| | | | | | 0 | 1 | 2 | 3 |
| Archaea (total) | 217 | 10372 | 1533 | 673 | 8402 | 1750 | 204 | 16 |
| Crenarchaeota | 60 | 2814 | 914 | 531 | 1553 | 1093 | 152 | 16 |
| Acidilobales | 2 | 92 | 16 | 9 | 67 | 25 | 0 | 0 |
| Desulfurococcales | 13 | 609 | 181 | 38 | 399 | 201 | 9 | 0 |
| Fervidicoccales | 1 | 45 | 8 | 7 | 33 | 9 | 3 | 0 |
| Sulfolobales | 29 | 1369 | 521 | 78 | 793 | 553 | 23 | 0 |
| Thermoproteales | 15 | 699 | 188 | 399 | 261 | 305 | 117 | 16 |
| Euryarchaeota | 144 | 6970 | 520 | 25 | 6432 | 531 | 7 | 0 |
| DHVE2 group | 2 | 92 | 31 | 0 | 61 | 31 | 0 | 0 |
| Archaeoglobales | 7 | 339 | 27 | 0 | 312 | 27 | 0 | 0 |
| Halobacteriales | 30 | 1481 | 106 | 0 | 1375 | 106 | 0 | 0 |
| Methanobacteriales | 13 | 559 | 35 | 0 | 524 | 35 | 0 | 0 |
| Methanocellales | 3 | 154 | 13 | 4 | 137 | 17 | 0 | 0 |
| Methanococcales | 16 | 605 | 32 | 0 | 573 | 32 | 0 | 0 |
| Methanomicrobiales | 9 | 449 | 26 | 0 | 423 | 26 | 0 | 0 |
| Methanopyrales | 1 | 35 | 8 | 2 | 27 | 6 | 2 | 0 |
| Methanosarcinales | 35 | 1958 | 172 | 11 | 1778 | 177 | 3 | 0 |
| Thermococcales | 20 | 929 | 40 | 0 | 889 | 40 | 0 | 0 |
| Methanomassiliicoccales | 3 | 138 | 13 | 6 | 120 | 17 | 1 | 0 |
| Thermoplasmatales | 5 | 231 | 17 | 1 | 213 | 18 | 0 | 0 |
| Korarchaeota | 1 | 46 | 3 | 3 | 41 | 4 | 1 | 0 |
| Nanoarchaeota | 1 | 44 | 3 | 1 | 40 | 4 | 0 | 0 |
| Thaumarchaeota | 11 | 498 | 93 | 107 | 337 | 122 | 39 | 0 |
| Cenarchaeales | 1 | 46 | 6 | 8 | 35 | 8 | 3 | 0 |
| Nitrosopumilales | 5 | 218 | 27 | 29 | 169 | 42 | 7 | 0 |
| Nitrososphaerales | 3 | 141 | 44 | 53 | 66 | 53 | 22 | 0 |
| unclassified | 2 | 93 | 16 | 17 | 67 | 19 | 7 | 0 |

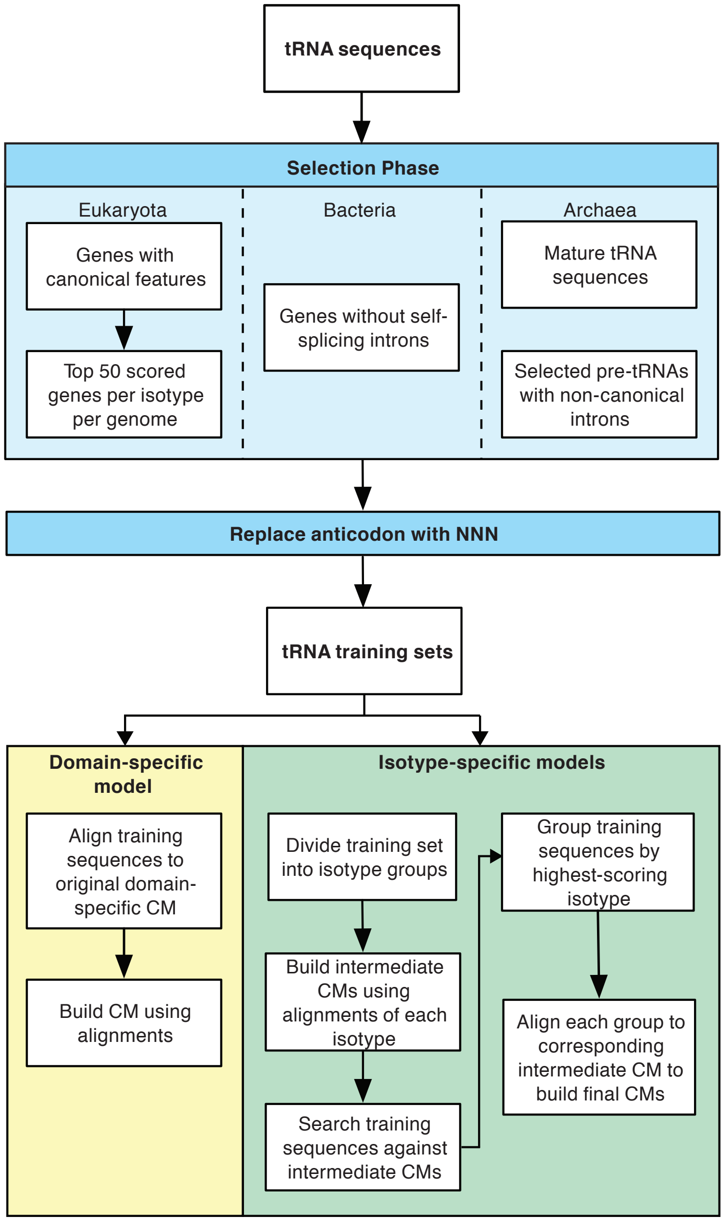


**Supplementary Figure S1.** Schematic diagram of covariance model generation. A training set for covariance model generation was created for each domain of life. Domain-specific CMs were built for eukaryotes, bacteria, and archaea with two rounds of training to build isotype-specific covariance models.

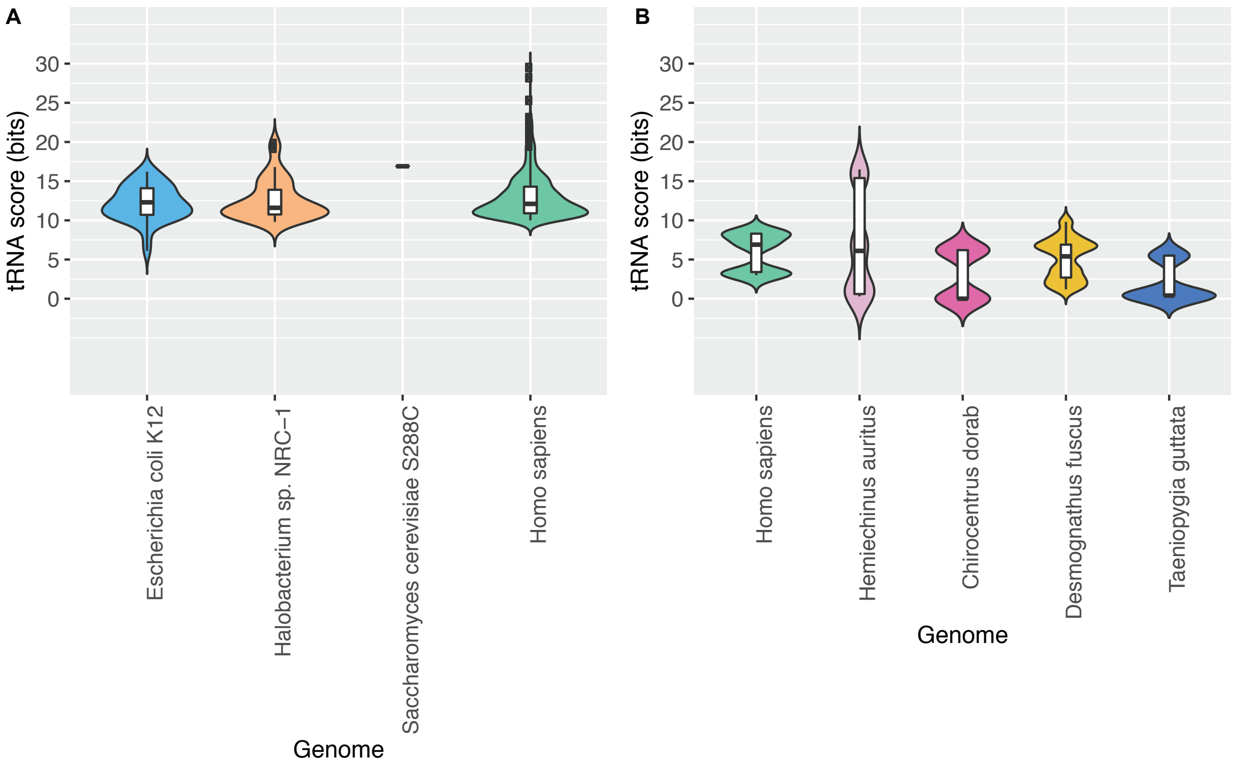


**Supplementary Figure S2.** tRNAscan-SE 2.0 score prediction distributions to assess false positives in (A) virtual nuclear genomes using corresponding domain search modes, and (B) virtual mitochondrial genomes using vertebrate covariance models. tRNA prediction results are also summarized in Supplementary Tables S3 and S4. Based on these distributions, a default cutoff of 20 bits was not changed from the original v1.3. The boxes inside the violin plot represent the interquartile range while the black bands represent the median of the tRNA scores.

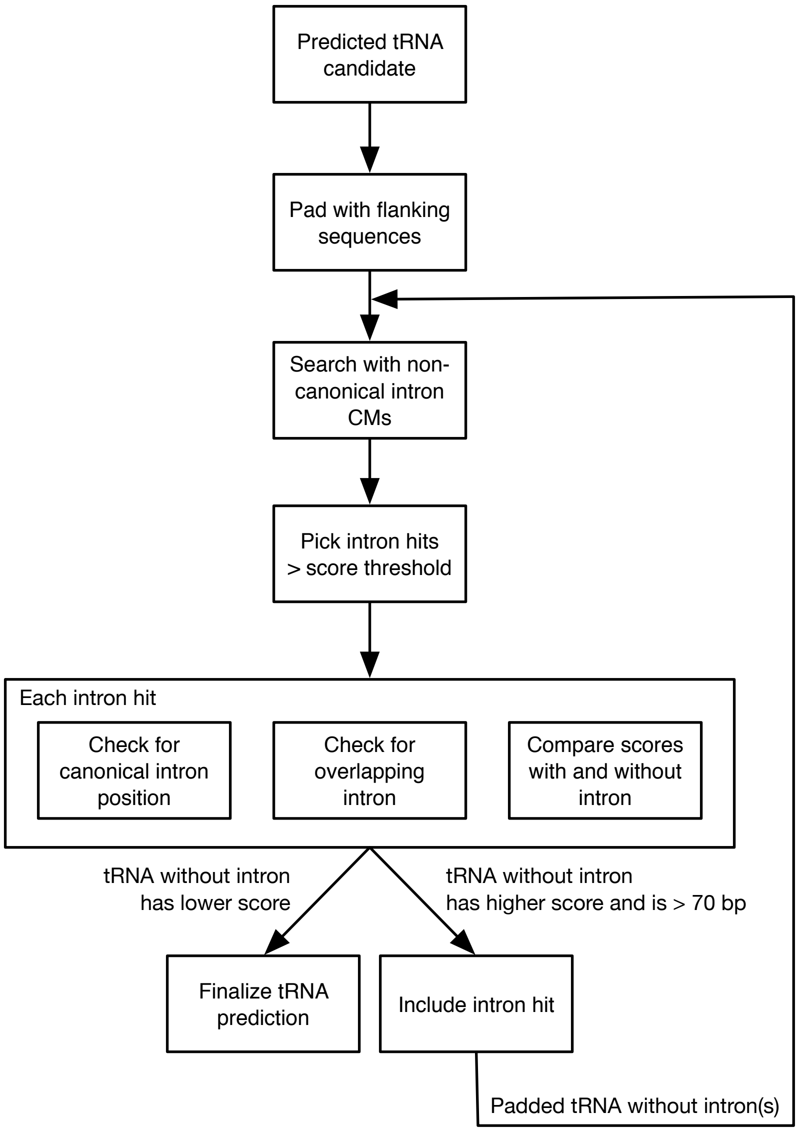


**Supplementary Figure S3.** Schematic diagram for detecting noncanonical introns in archaeal tRNA genes. The tRNA gene candidates were padded with flanking sequences and scanned with covariance models that were trained with previously identified noncanonical introns in archaeal tRNAs. The detected introns were characterized by their scores and positions. An iterative process was used to enable detection of multiple introns that are located in close proximity and requires the removal of one intron at a time, resulting in formation of the bulge-helix-bulge secondary structure motif of any potentially remaining intron(s).

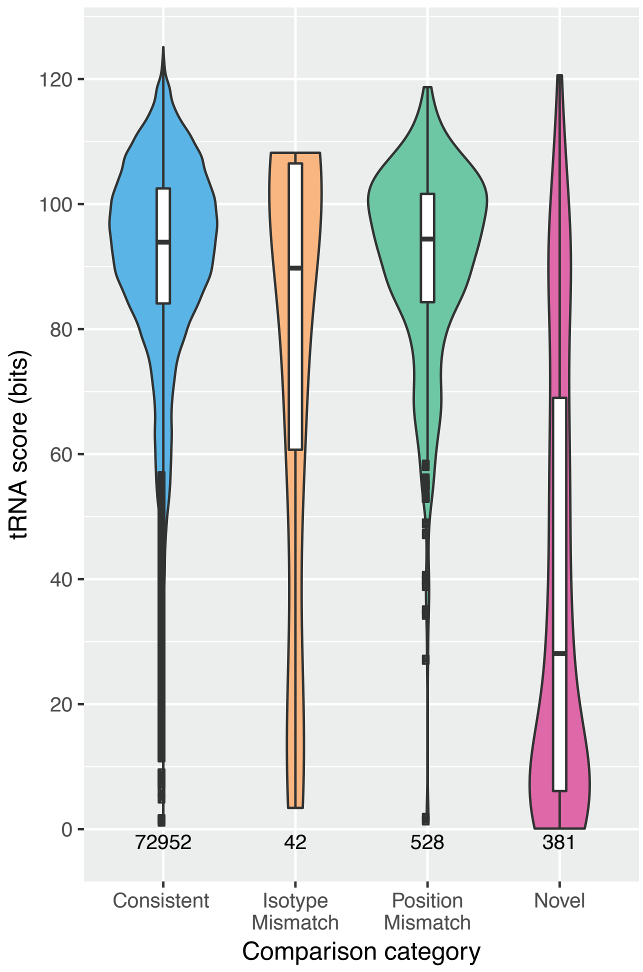


**Supplementary Figure S4.** Mitochondrial tRNA prediction comparison with NCBI RefSeq (7). 73,903 mt-tRNA genes in 3,345 mitochondrial genomes were predicted using tRNAscan-SE 2.0 with vertebrate mitochondrial search mode and score cutoff of 0 bits. Predictions were compared with 73,674 NCBI RefSeq gene annotations and were grouped into four categories: (1) Consistent – predicted genes are consistent with RefSeq annotations, (2) Isotype Mismatch – predicted genes and RefSeq annotations have different isotype classification, (3) Position Mismatch – predicted genes and RefSeq annotations are on different strands or have different start/end positions but do overlap, and (4) Novel – predicted genes are not annotated in RefSeq. The number below the plot of each category represents the total number of predicted genes for the specified category. The boxes inside the violin plot represent the interquartile range while the black bands represent the median of the tRNA scores.

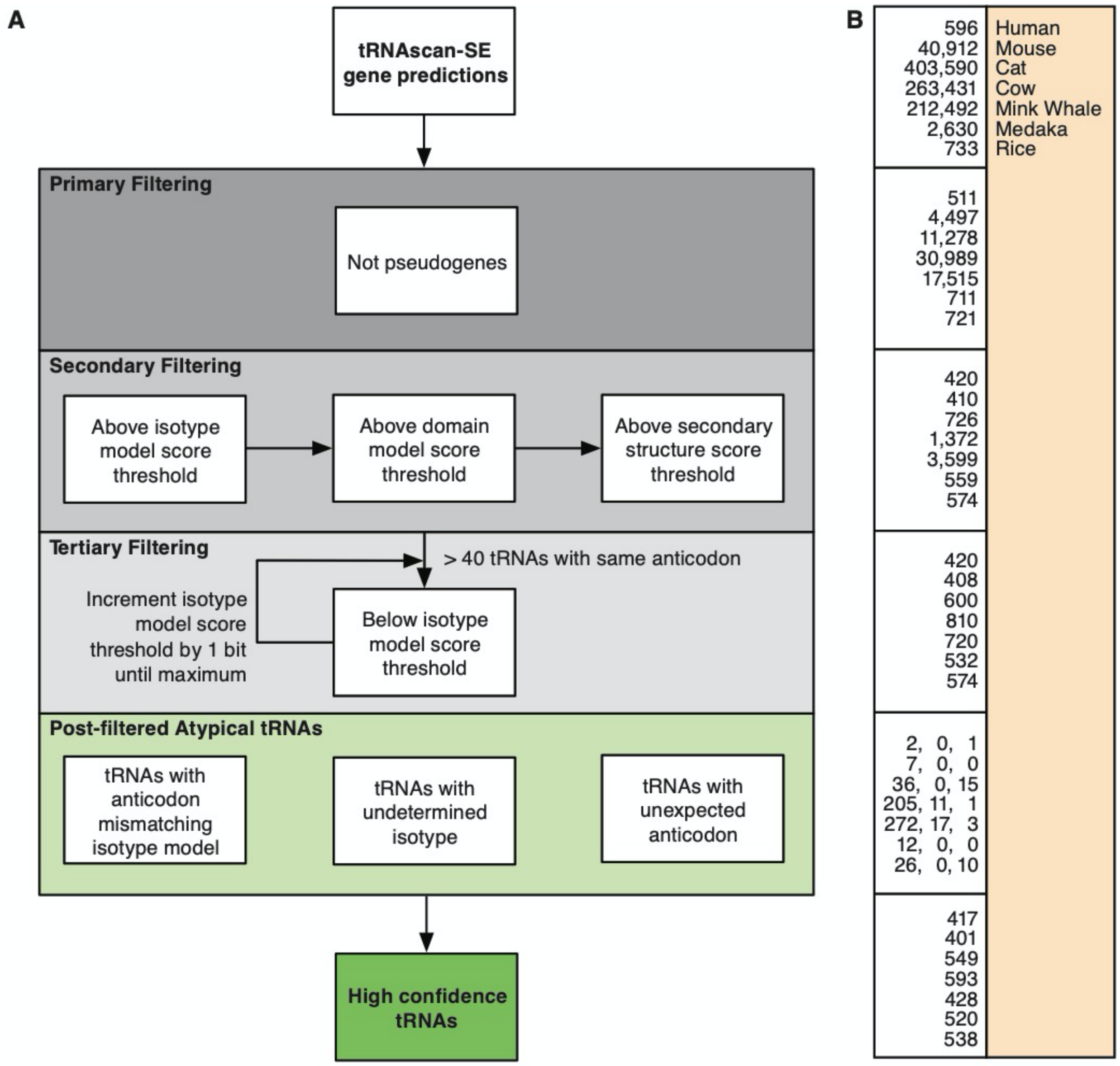


**Supplementary Figure S5.** Schematic diagram of post-filtering process for identifying high confidence tRNA genes. (A) Flowchart represents the filtering steps used for tRNA predictions in large eukaryotes to better distinguish between tRNA-derived SINEs and real tRNA genes. (B) Counts of tRNA gene predictions of selected genomes are listed at each filtering step.

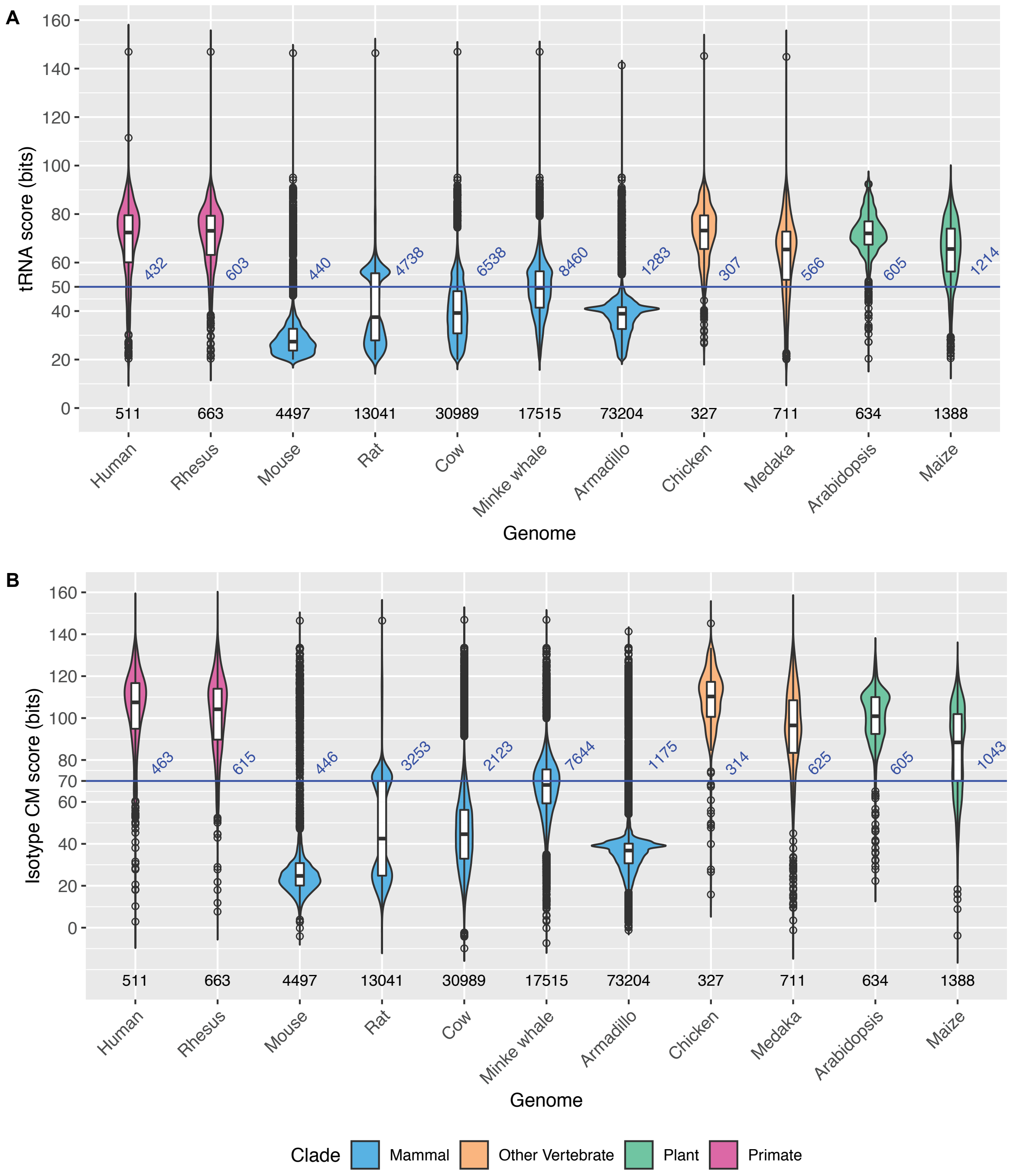

A
tRNA score (bits)
160
140
120
100
80
60
50
40
20
0
432
603
440
4738
6538
8460
1283
307
566
605
1214
511
663
4497
13041
30989
17515
73204
327
711
634
1388
Human
Rhesus
Mouse
Rat
Cow
Minke whale
Armadillo
Chicken
Medaka
Arabidopsis
Maize
Genome
B
Isotype CM score (bits)
160
140
120
100
80
70
60
40
20
0
463
615
446
3253
2123
7644
1175
314
625
605
1043
511
663
4497
13041
30989
17515
73204
327
711
634
1388
Human
Rhesus
Mouse
Rat
Cow
Minke whale
Armadillo
Chicken
Medaka
Arabidopsis
Maize
Genome
Clade
Mammal
Other Vertebrate
Plant
Primate

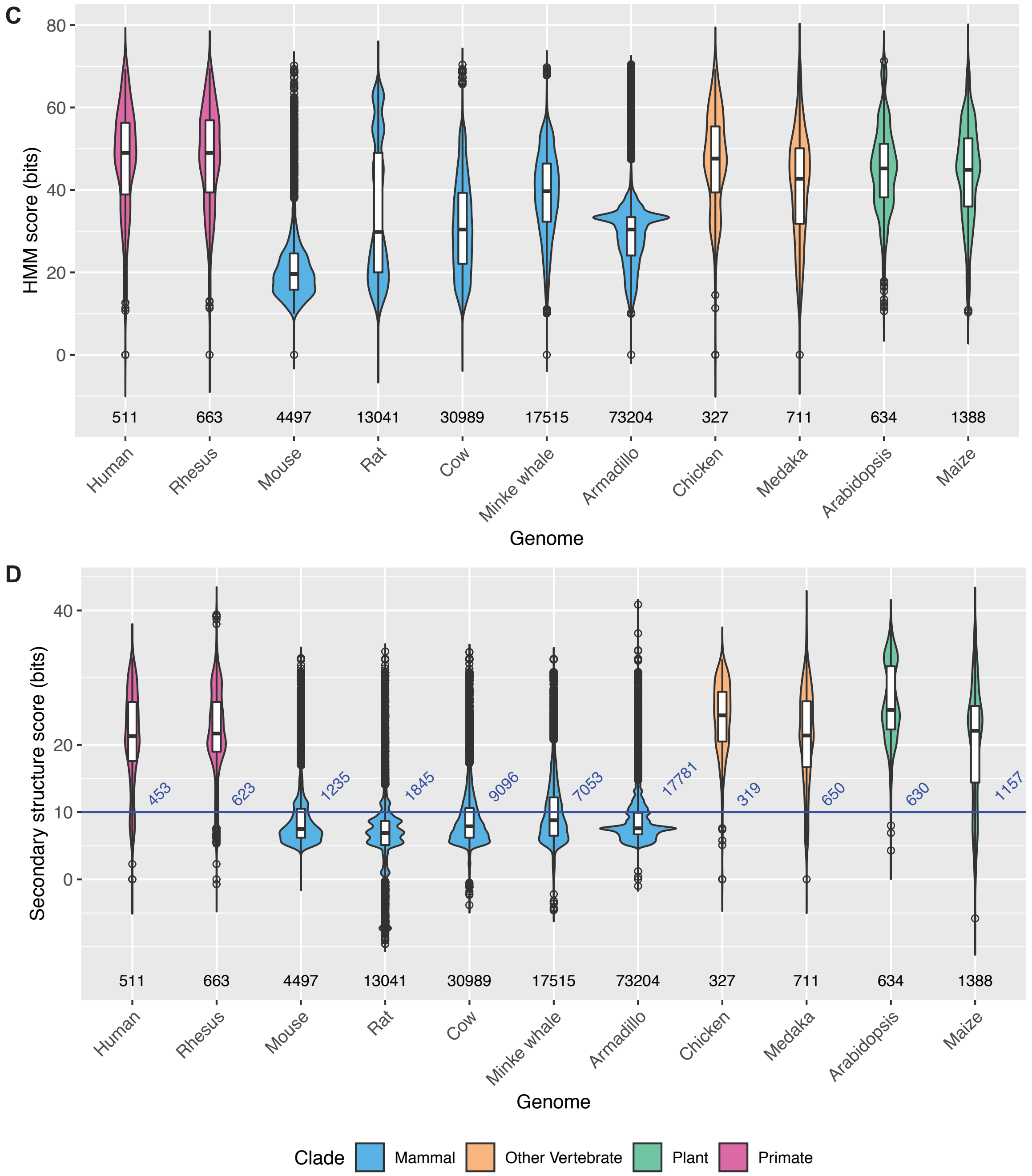


**Supplementary Figure S6.** tRNA prediction score distributions of selected large eukaryotic genomes, showing a wide range of counts and scores. Predictions classified as pseudogenes have been excluded. Distributions of four score types are included: (A) tRNA (domain-specific) scores, (B) isotype-specific scores, (C) HMM scores (scores based on primary sequences), and (D) secondary structure scores. The horizontal blue lines represent score thresholds used in the post-filtering approach for identifying high confidence tRNA genes. No single filtering threshold is sufficient for removing tRNA-derived repetitive elements for all species, thus a multi-stage filter was developed. The numbers below the violin plots of each genome are the total number of tRNA predictions represented in the distributions. The numbers at the horizontal blue lines are the number of tRNA predictions at the corresponding score thresholds. The boxes inside the violin plot represent the interquartile range while the black bands represent the median of the tRNA scores.

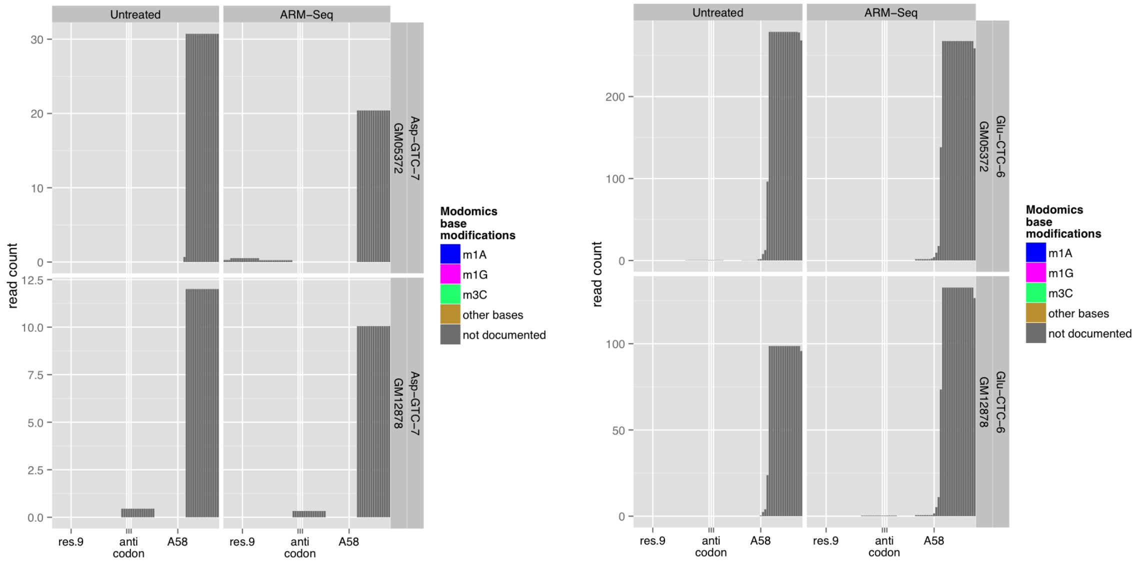


**Supplementary Figure S7.** Read coverage of tRNA-Asp-GTC-7 and tRNA-Glu-CTC-6 in human from ARM-Seq study (4). Sequencing reads only aligned to the 3' end of the transcripts.

**A**

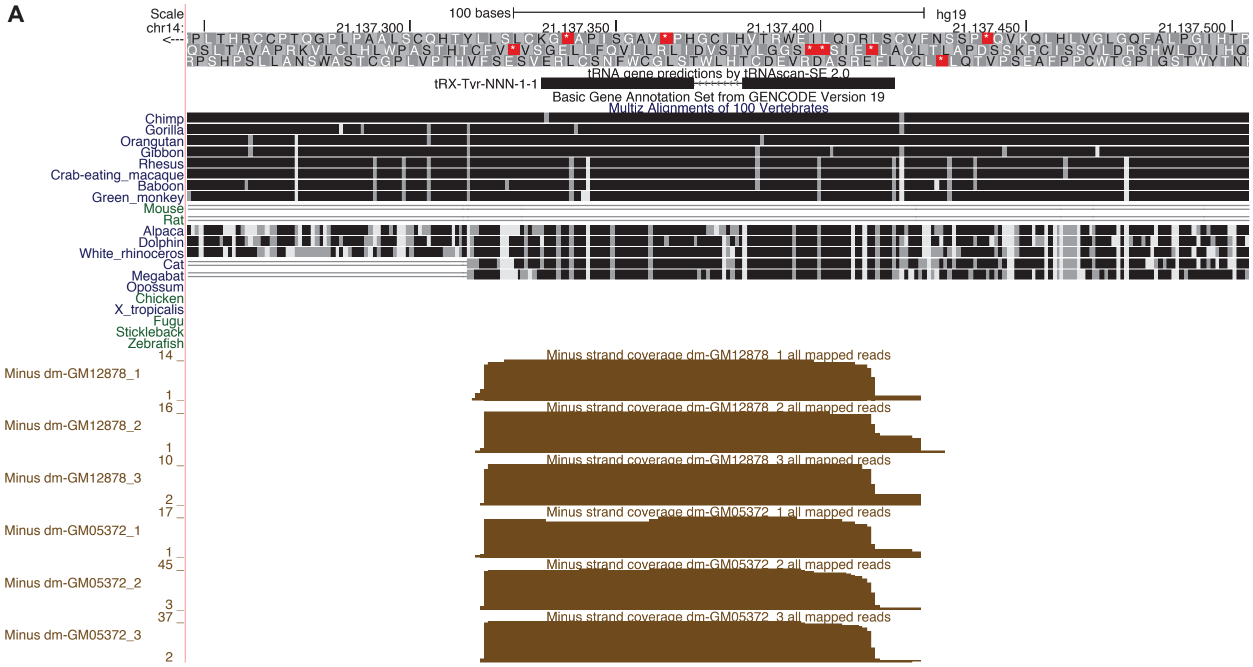


**B**

| Genome | GtRNAdb Gene Symbol | Score (bits) | Feature |
|---|---|---|---|
| Human | tRX-Tyr-NNN-1-1 | 22.8 | Uncertain function |
| Chimp | N/A | 19.1 | Below default score cutoff |
| Gorilla | tRX-Tyr-NNN-3-1 | 20.7 | Uncertain function |
| Rhesus | tRX-Tyr-NNN-4-1 | 39.5 | Uncertain function |
| Baboon | tRX-Tyr-NNN-4-1 | 39.5 | Uncertain function |
| Alpaca | tRNA-Tyr-GTA-3-6 | 75.4 | High confidence |
| Dolphin | tRNA-Tyr-GTA-6-1 | 72.9 | High confidence |
| White rhinoceros | tRNA-Tyr-GTA-3-3 | 75.4 | High confidence |
| Cat | tRNA-Tyr-GTA-3-2 | 75.4 | High confidence |

**Supplementary Figure S8.** tRX-Tyr-NNN-1-1 in human shows uniform expression as a precursor tRNA. (A) The UCSC Genome Browser (8) visualization shows tRX-Tyr-NNN-1-1, a predicted tRNA gene candidate for decoding tyrosine with anticodon GUA. The predicted gene has a 12-nt intron represented by the arrowed portion of the track item and is conserved across primates and some other mammals. The ARM-Seq coverage tracks show uniform coverage across the gene body and the intron instead of higher coverage at the mature sequence region, indicating that the gene transcript may not be processed as a typical tRNA. (B) Orthologs in mammals based on multi-genome alignments are listed with GtRNAdb gene symbols, scores, and feature classification.

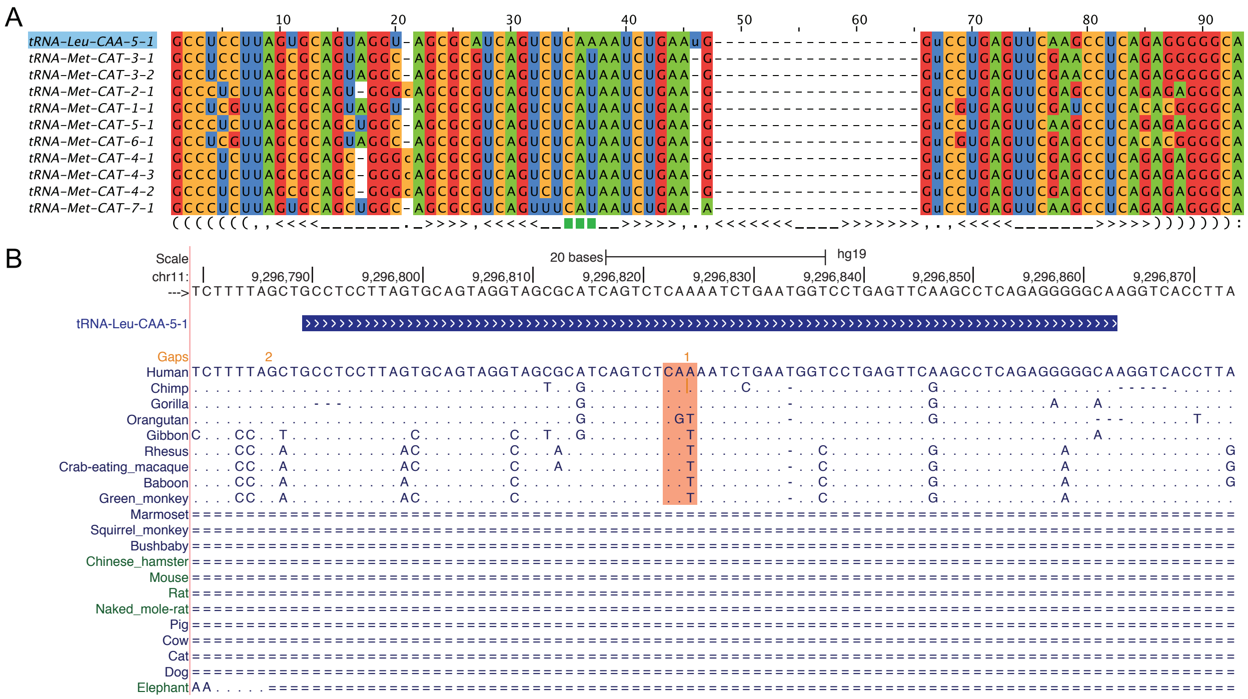


**Supplementary Figure S9.** Sequence conservation of tRNA-Leu-CAA-5-1 in human. (A) Alignment of tRNA-Leu-CAA-5-1 with all predicted $tRNA^{Met}$ sequences from human shows high conservation supporting its ancestry as a $tRNA^{Met}$ gene. (B) UCSC Genome Browser view of tRNA-Leu-CAA-5-1 locus that is shared with other primates, but not other mammals ("="). Anticodon CAA (highlighted) is conserved in human, chimp, and gorilla (identity with human reference sequence shown with "."). The genomes of other primates have anticodon CAT (corresponding to $tRNA^{Met}$) of this gene instead.
